# Detection of Region-specific Fiber Damage within Injured Spinal Cord Using Advanced Diffusion MRI

**DOI:** 10.1101/2025.07.02.662840

**Authors:** Feng Wang, Junzhong Xu, Tung-Lin Wu, Pai-Feng Yang, Li Min Chen, John C. Gore

## Abstract

This study aimed to evaluate diffusion parameters derived from diffusion tensor imaging (DTI) and spherical mean technique (SMT) for detecting region-specific, fine-grained tissue damage and white matter (WM) tract disruptions following spinal cord injury (SCI). Diffusion MRI data were acquired from the cervical spinal cord of monkeys before and after a unilateral dorsal column lesion at the C5 level, using a 9.4T scanner. Parametric maps derived from DTI and SMT effectively detected regional fiber damage around 16 weeks post-injury. Post-mortem silver staining served as the ground truth for assessing region-specific fiber damage. Diffusion MRI maps aligned well with histological measures and captured the severity of WM damage at the lesioned segment (in an order of dorsal > ventral > lateral WM tracts) and along the dorsal column tract across segments (in an order of lesion center > rostral > caudal). Among the diffusion parameters, fractional anisotropy (FA), axonal volume fraction (V_ax_), radial diffusivity (RD), and extra axonal transverse diffusivity (D_ex_) showed most significant changes at and around the lesion site where severe tissue damage occurred. FA, V_ax_, and axial diffusivity (AD) exhibited marked changes in dorsal column proximal to the lesion center, where moderate axonal damage occurred. Additionally, AD and FA showed the greatest sensitivity (true positive rate) and specificity (true negative rate) to mild fiber disruption and demyelination in regions distal to the lesion. Overall, FA provided the highest sensitivity and specificity for detecting fiber degeneration and demyelination, while V_ax_ demonstrated the strongest spatial correlation with histologic markers of regional fiber damage. The combination of DTI and SMT thus offers reliable biomarkers for assessing SCI.

## 1 INTRODUCTION

Traumatic spinal cord injury (SCI) disrupts the structural integrity and normal functions of the spinal cord, leading to sensory, motor, and autonomic functional deficits. The injury can induce cell death, demyelination, and axonal loss,^1, 2^ as well as edema and the formation of cysts^3^ in spinal cord tissues over time. Non-invasive magnetic resonance imaging (MRI) is well-suited for monitoring the progression of damage and the recovery of injured spinal cord tissues.^4–7^ MRI metrics provide comprehensive insights into structural, functional, and molecular changes following injury,^4–10^ which may serve as mechanism-informative biomarkers for aiding in the screening of therapeutic interventions in preclinical animal models and evaluating the treatment effects in clinical trials.

Diffusion MRI is sensitive to detecting and assessing disruptions in tissue microstructure and cellular integrity, providing valuable insights into the degeneration and repair processes within injured spinal cords over time.^4, 11, 12^ Currently, diffusion tensor imaging (DTI) is the most widely used technique due to its relatively high reliability and ease of data acquisition. DTI uses a three-dimensional tensor to model the anisotropy of water movements in white matter (WM), yielding metrics that are highly influenced by cell density, microstructure, and fiber density.^13^ DTI-derived fractional anisotropy (FA), mean diffusivity (MD), axial diffusivity (AD), and radial diffusivity (RD) are sensitive to tissue properties such as axonal density and myelination level.^8, 9^ However, DTI has two notable limitations. Firstly, it assumes a Gaussian signal phase distribution, which becomes unreliable at high *b*-values in structured tissues like the spinal cord. Secondly, it uses a single averaged tensor to describe water compartment contributions, reducing its specificity in capturing complex tissue microstructure.

To overcome these limitations, more advanced biophysical models have been developed to describe the signals measured in diffusion MRI. Some studies have applied multi-compartmental diffusion imaging in the spinal cord.^14–18^ The spherical mean technique (SMT) models the microscopic diffusion process, enabling the mapping of neurite density and compartment diffusivities.^16, 17^ SMT may offer potentially significant advantages for assessing fiber loss. By taking the spherical mean of diffusion signals across various gradient directions for a given *b*-value, it provides orientation-invariant indices, including the apparent axonal volume fraction (V_ax_), intrinsic diffusivity (D_ax_), and extra-axonal transverse diffusivity (D_ex_). In practice, applying advanced diffusion models to the spinal cord poses challenges due to the low signal-to-noise ratio (SNR) inherent in high-resolution imaging and the impact of physiological noise. By averaging all diffusion-weighted images over a given *b*-value, SMT inherently increases the SNR, potentially enhancing the robustness of multi-compartmental model fitting.^18^ Additionally, while the normal spinal cord is predominantly composed of WM fibers aligned in the same direction, other nerve structures, such as dorsal and ventral roots, are oriented approximately perpendicular to the main cord direction. Moreover, posture variations, such as relaxation or stretching, can cause spinal cord undulation, which may affect diffusion tensor imaging (DTI) measurements.^19^ SCI induces neurodegeneration that can alter the organization of nerve fibers, such as increasing the orientation dispersion of WM fibers.^15^ Since the SMT model is insensitive to fiber orientation and organization, it may provide a complementary, straightforward, and reliable measure for characterizing fiber degeneration and loss following SCI.

Axonal demyelination, degeneration, and loss are primary pathological processes led to irreversible neurological impairment in subjects with traumatic SCI. Among preclinical models, the unilateral dorsal column lesion (DCL) is particularly well-suited for evaluating diffusion MRI’s ability to detect focal, spatially restricted white matter (WM) damage because it is designed to disrupt ascending somatic afferents to the brain without affecting other white matter tracts and grey matter regions.^20–23^ Unlike previous longitudinal studies using this model,^4, 11, 12^ this study focused on comparing DTI- and SMT-derived metrics for detecting fine-scale microstructural damage in specific WM tracts and gray matter (GM) regions at the lesion site, as well as in segments proximal and distal to the injury along the cord. We assessed the sensitivity (true positive rate) and specificity (true negative rate) of various diffusion metrics in detecting tissue damage across severity levels and evaluated their performance by using histological measures of region-specific mild, moderate, and severe fiber loss as ground truth. Furthermore, this study aimed to determine whether parameters derived from SMT outperform those derived from DTI in detecting regionally specific fiber damage.

## 2 MATERIAL AND METHODS

### 2.1 General information and SCI model - dorsal column transection

Ten male adult squirrel monkeys (*Saimiri sciureus*) were studied, and seven of them underwent a unilateral dorsal column transection at cervical level 5 (C5). This DCL SCI model is designed to selectively disrupt the ascending dorsal column tract/pathway (DP). Details of the surgical procedures can be found in Supporting Information and previous publications.^21, 24, 25^ All procedures were approved by the Institutional Animal Care and Use Committee (IACUC) at Vanderbilt University and adhered to the NIH guidelines for the care and use of laboratory animals.

### 2.2 *In vivo* MRI data acquisition

MRI data were acquired from the cervical spinal cords of ten monkeys before injury and seven monkeys around week 16 post-injury. During MRI data acquisitions, each monkey was anesthetized with isoflurane (0.8-1.2%), delivered in a 70:30 N_2_O/O_2_ mixture, and mechanically ventilated. Vital physiological signs were continuously monitored and maintained at stable levels throughout the imaging session.

All MR images were acquired at 9.4T using a customized saddle-shaped transmit-receive quadrature surface coil^26^ (each of the two loops is 30-mm wide and 30-mm long, mounted on 45-mm-diameter cylindrical surface) positioned around monkey’s neck. The coil placement was confined to the cervical spinal cord, which reduced unwanted signals from distant moving tissues and increased the signal-to-noise ratio. Magnetization transfer contrast (MTC)-weighted structural images were acquired in multiple orientations with enhanced contrast (Supporting Information),^4^ facilitating the precise placement of diffusion-weighted imaging planes around the lesion center (Fig. 1A).

**Figure 1.**
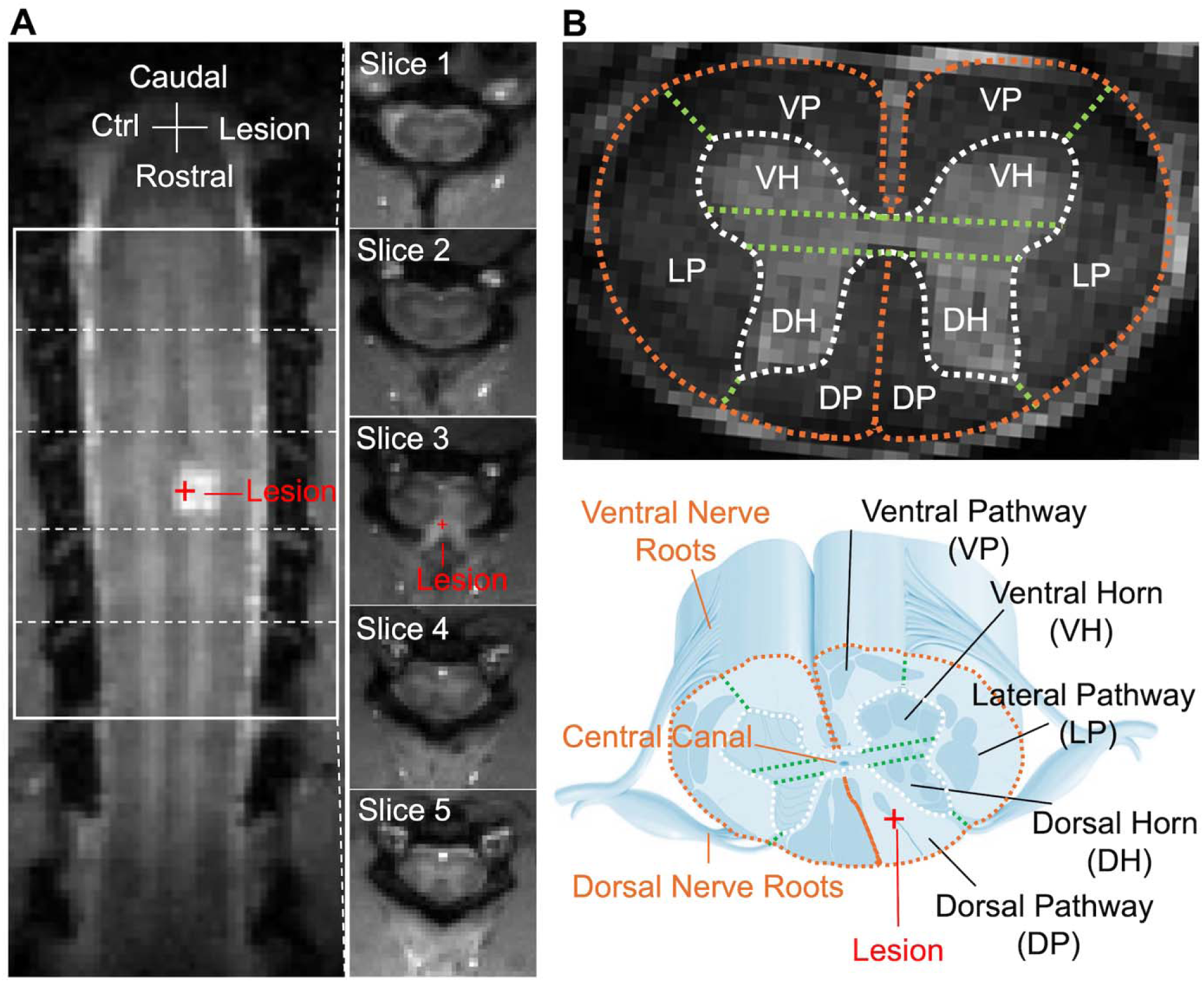
Placement of the diffusion MRI and regions of interest (ROIs). (A) Horizontal and axial MRI image with magnetization transfer contrast (MTC) identified the lesion as a hyperintensity region at the dorsal column on one side of the cervical spinal cord at the C5 level in one representative monkey. The white boxes on the horizontal image (left) indicate the placements of five axial image slices over the cervical enlargement segments for acquiring diffusion MRI data. Red cross indicates the location of the lesion site at axial slice 3 level. (B) Selected corresponding ROIs on high-resolution MTC image based on schematic diagram showing the organization of the spinal cord. The landmarks such as central canal, ventral nerve roots, and dorsal nerve roots are used for ROI selection for white matter regions (VP: ventral pathway, LP: lateral pathway, DP: dorsal pathway) and gray matter regions (DH: dorsal horn, VH: ventral horn). DH ROIs are below the central canal level and VH ROIs above the central canal level. The boundaries are indicated by dashed lines.

The diffusion-weighted spin-echo sequence used an echo planar imaging readout (TR/TE = 3000/33 ms, 4 shots, resolution = 0.333x0.333 mm^2^, slice thickness = 3 mm, 5 slices). Three *b*-shells were acquired with *b* values of 750, 1000, and 2000 s/mm^2^, sampling 30 directions per shell (equally spaced). At least three non-diffusion-weighted scans were acquired, each preceding the acquisition of a *b*-value shell. Multi-shot EPI was used to mitigate distortion and local signal dropouts. Navigator and phase correction were applied to minimize motion artifacts and ghosting. An extra navigator echo per readout train was collected without phase encoding prior to the acquisition of the actual image data. This approach improved signal stability in segmented multi-shot EPI, corrected for motion-induced phase variations, and addressed frequency drift issues. To further reduce motion artifacts, we synchronized the diffusion data acquisition (TR = 3000 ms) with the respiration cycle (1500 ms/cycle). No evident artifacts were observed in diffusion-weighted images (*Sup. Fig. S1*). High-resolution MTC and T_2_-weighted (T2W) anatomical images were acquired at 0.125 x 0.125 mm^2^ and 0.250 x 0.250 mm^2^ respectively, using the same geometry (Supporting Information). The total acquisition time for both anatomical and diffusion MRI data was approximately 32 minutes.

### 2.3 MRI Data Analyses

MRI data were analyzed using MATLAB (The MathWorks). V_ax_, D_ax_, and D_ex_ were derived from SMT using custom MATLAB code.^16^ Conventional DTI parameters, including FA, AD, RD, and MD, were quantified using the DTI Toolbox^27^ in MATLAB based on diffusion data from the *b* = 1000 s/mm^2^ shell.

The non-diffusion-weighted SE-EPI (A_0_) images were diffeomorphically registered slice-wise to the high-resolution MTC images, and all other diffusion weighted volumes (A_1_-A_30_) were registered to A_0_ image using a rigid registration algorithm (Sup. Fig. S1). MTC, T_2_-weighted (T2W), and diffusion-weighted SE-EPI images effectively differentiate normal WM and GM with good contrast (Fig. 1 and Sup. Fig. S1), facilitating registration and further regional analysis.

### 2.4 Histology of post-mortem spinal cord tissue

Histological silver staining, which is used to selectively visualize and assess axonal integrity (highlighting intact axons), myelin and demyelination, as well as neurodegeneration, was performed on post-mortem spinal cord tissue in all 10 subjects, one or two weeks after the final MRI scan (Supporting Information).^5, 28^ Histology data from six SCI animals with transverse tissue sections were quantified and used to validate the diffusion MRI results.

The histological RGB image was inverted and converted to the CIELAB color space (defined by the International Commission on Illumination in 1976) using MATLAB, where Lightness (0-100 from black to white), a measure that is similar to intensity, is used for assessing fiber density. The regional mean of Lightness was quantified and served as histological inD_ex_ (HI) for validation of fiber density in each region of interest (ROI).

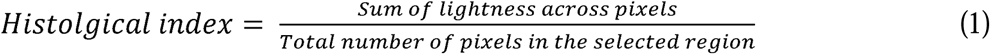

For each ROI, the histological measure was calculated by averaging values from three sections, corresponding to a single MRI slice, and aligned with respective ROI values from respective MRI segment for comparison.

### 2.5 ROIs defined based on MRI and histological landmarks and boundaries

High-resolution MTC and respective histological images were used as references for the manual selection of ROIs for quantification, based on the known anatomy and landmarks of monkey cervical spinal cord (Fig. 1B and Sup. Fig. S2).^29^ The ROIs on each side of a single spinal cord segment were manually delineated along the regional boundaries using MATLAB, with voxels along the border excluded from the analysis to minimize partial volume effects (Fig. 1 and Sup. Fig. S2).

The spinal cord is bilaterally symmetrical, and 5 ROIs were defined for each side of spinal cord, resulting a total of 10 ROIs for each segment (Fig. 1 and Sup. Fig. S2). Three WM tracts and two GM ROIs were defined based on histological characteristics identified from fiber stains and commonly recognized anatomical regions on the atlas in monkeys.^29^ The landmarks and boundaries are shown in a schematic of spinal cord GM regions and WM tracts and ROIs are indicated on a histological fiber stain and MTC image (Fig. 1B and Supporting Fig. S2). The spinal cord WM was segmented into dorsal, lateral, and ventral pathways (DP, LP, and VP, respectively), while the GM was parcellated into the dorsal and ventral horns (DH and VH, respectively) on each side of spinal cord.

### 2.6 Statistical Analysis

A two-sided Wilcoxon rank sum test was performed to evaluate the statistical significance of differences in diffusion MRI measures (n =10 healthy, n = 7 injured) and histological measure (n = 10 healthy, n = 6 injured) for each individual ROI, comparing lesion and contralateral non-lesion side with healthy spinal cord tissues. A *p*-value < 0.05 was considered statistically significant. Because the high prevalence rate of this targeted injury model requested small sample size for ROC analysis, the true positive rate (TPR), true negative rate (TNR), and area under the ROC curve (AUC) were obtained for assessing sensitivity, specificity, and diagnostic performance respectively (Supporting Information). For each measure, values from healthy tissue ROIs were used as the gold standard for comparison. The strength of the linear relationship between different MRI and histological measures was determined using the Pearson correlation coefficient.

## 3 RESULTS

### 3.1 Diffusion MRI detect and differentiate tissue property changes in specific ROIs at and around the lesion site

WM ROIs displayed much lower intensity than GM ROIs on high-resolution MTC images (Fig. 1B), reflecting the high density of myelinated axons in WM. Both DTI- and SMT-derived parametric maps (Fig. 2) detected tissue property changes at the lesion center on the injury side of the spinal cord (slice 3 in Fig. 1). Representative diffusion parametric maps delineate normal WM and gray matter (GM) in the pre-lesion spinal cord (Fig. 2A). Although both D_ax_ and D_ex_ maps showed increased values at the lesion site (Fig. 2B), the D_ex_ map exhibited a much larger signal change on the lesion side than D_ax_. Similarly, the RD map showed a larger increase at the lesion center compared to AD. A drastic decrease in V_ax_ was observed at the lesion center, concurrent with a reduction in FA (Fig. 2B). These observations were generally consistent for all animals,^30^ despite variations in injury severity among subjects.

**Figure 2.**
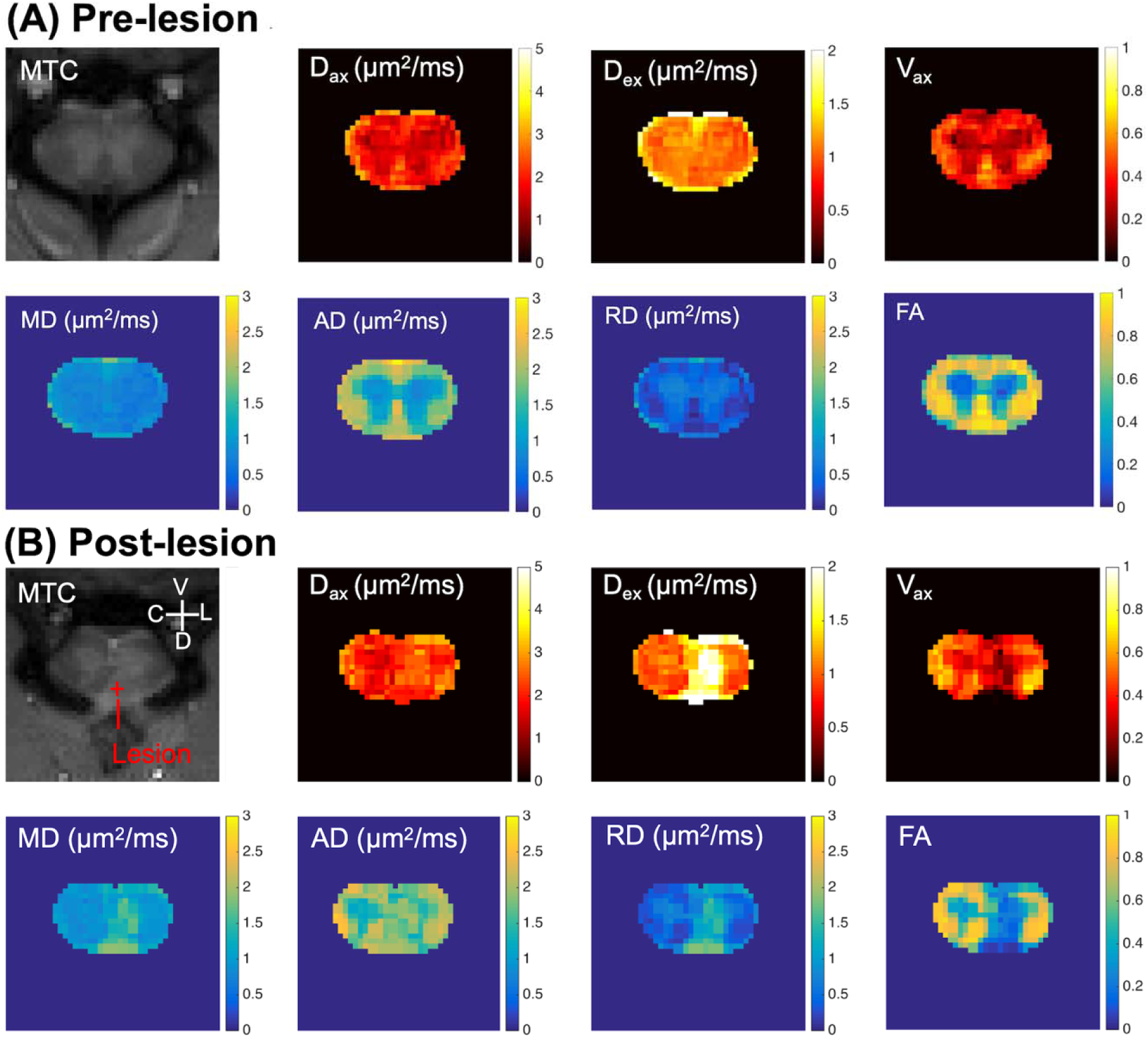
Comparison of diffusion parametric maps of the center slice from one representative subject. (A) Pre-lesion. (B) Post-lesion. MTC, magnetization transfer contrast. Measures from spherical mean technique: D_ax_, intrinsic axonal diffusivity; D_ex_, extra axonal transverse diffusivity; V_ax_, apparent axonal volume fraction. Measures from diffusion tensor imaging: MD, mean diffusivity; AD, axial diffusivity; RD, radial diffusivity; FA, fractional anisotropy. Red cross indicates the column lesion on one side of dorsal pathway (DP). V, ventral; D, dorsal; C, non-lesion contralateral side, L, lesion side. The segment directly underwent unilateral dorsal column lesion is shown.

The decrease in V_ax_ and FA extended both rostrally and caudally along the dorsal column tract, reaching even the distal segments in the rostral direction (Fig. 3). Additionally, the rostral region proximal to the lesion exhibited more severe tissue disruption than the caudal region (Fig. 3). While AD and D_ax_ decreased in some segments distal to the lesion center (indicated by white asterisks in Fig. 3), RD and D_ex_ did not detect changes in tissue properties in these distal segments. Unlike other diffusion metrics, both AD and D_ax_ exhibited directionally different changes in the dorsal column depending on the severity of axonal damage (indicated by red and white asterisks in Fig. 3).

**Figure 3.**
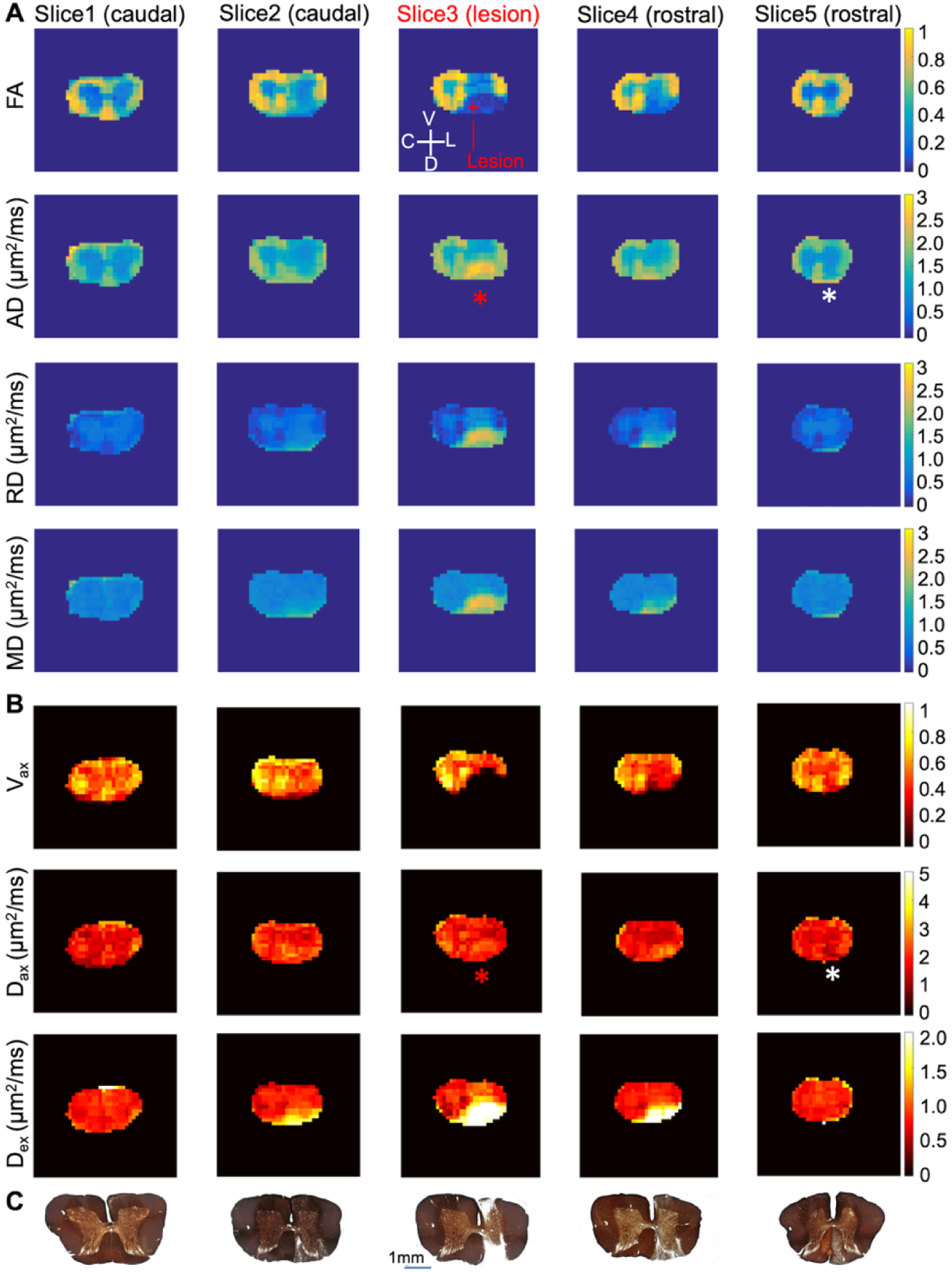
Comparison of diffusion parametric maps and corresponding histological maps in one representative spinal cord with a unilateral dorsal column lesion across five axial slices from caudal to rostral along the cord. (A) DTI-derived parameter maps. (B) SMT-derived parameter map. (C) Silver staining of corresponding spinal cord tissue. V: ventral; D: dorsal; C: non-lesion contralateral side; L: lesion side. FA, fractional anisotropy; AD, axial diffusivity; RD, radial diffusivity; MD, mean diffusivity. V_ax_, apparent axonal volume fraction; D_ax_, intrinsic axonal diffusivity; D_ex_, extra axonal transverse diffusivity. The red cross indicates the lesion site. The red asterisks indicate the increase of AD and D_ax_ at the lesion site while the white asterisks indicate unilateral decrease in slice distal to the lesion on the rostral side. Fonts are too small.

### 3.2 Spatially constrained tissue damage and histological validation

Silver staining confirmed myelin loss, axon degeneration and tissue damage in the injured spinal cord, particularly around the targeted lesioned dorsal column pathway (DP), as well as cyst formation at the lesioned spinal segment (Fig. 3C). The presence of cyst formation at and around the lesion site, which was primarily composed of scar tissues, has also been confirmed in some subjects through additional histological staining, as reported in previous studies.^4, 5, 30^ Silver-stained sections further validated that reductions in V_ax_ and FA values were spatially aligned with regional demyelination and axonal damage in WM, both at and around the injury site (Fig. 3C). Severe damage was evident at the lesion site of the cord, especially in the DP and dorsal horn (DH). Proximal to the lesion site, the rostral segment showed more pronounced tissue damage than the caudal segment in this subject (Fig. 3C), a finding consistent with the expected disruption of ascending fibers on the rostral segments to the injury, which carries somatic sensory inputs from the spinal cord to the brain. Along the DP, the damaged regions extended beyond 3 mm in length (the length of one spinal segment) on both the caudal and rostral segments to the lesion site, reaching distal segments on the rostral side.

### 3.3 Comparison of regional diffusion characteristics between healthy and injured spinal cords

To determine the spatial distribution of tissue damage for this injury model, we performed an ROI-based group analysis. For each of the five ROIs (VH, DH, LP, VP, and DP), regional mean values of diffusion measures were compared between the injured spinal cords and healthy controls. Specifically, comparisons were made for the injured segment on the contralateral (non-lesion) side (slice 3 only) and for five segments on the lesion side (slices 1–5) (Figs. 4–5). On the lesion side, ROIs were analyzed in the injured segment (slice 3), two rostral segments proximal and distal to the lesion (slices 4 and 5, respectively), and two caudal segments proximal and distal to the lesion (slices 2 and 1, respectively). No diffusion metrics showed significant changes in the five selected ROIs on the contralateral side (Figs. 4–5). On the lesion side, FA and V_ax_ exhibited regionally significant decreases, with intra-segment variations from dorsal to ventral regions (WM: DP > VP > LP; GM: DH > VH) and inter-segmental changes along the DP, following the pattern: lesion center > rostral segment proximal to lesion > caudal segment proximal to lesion > rostral segment distal to lesion > caudal segment distal to lesion. Notably, AD and D_ax_ did not show significant changes in the DP at the lesion center across subjects; however, they decreased significantly in the DP of either the rostral or caudal segments (Figs. 4-5). In contrast, RD and D_ex_ exhibited greater increases at the lesion center compared to the adjacent rostral and caudal segments (Figs. 4-5).

**Figure 4.**
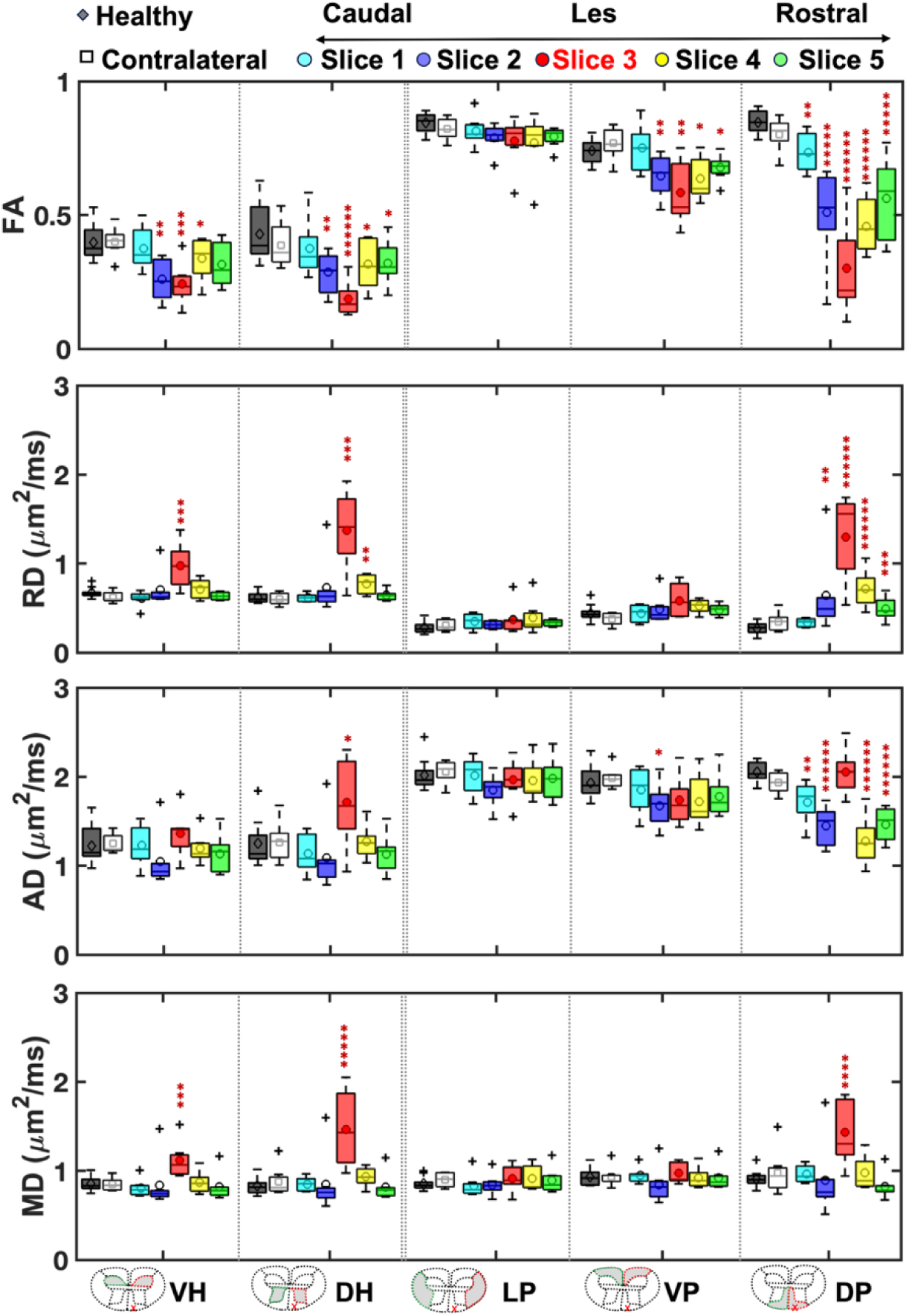
Regional comparison of DTI-derived diffusion measures in five ROIs across slices of injured spinal cords with those from healthy cords. Boxplots of DTI-derived FA, RD, AD, and MD metrics show the differences for the selected ROIs from the healthy spinal cords without injury, non-lesioned contralateral side of the lesioned segment (slice 3 as indicated in Figure 3), and different segments on the lesion side from caudal to rostral (slices 1-5 as indicated in Figure 3) of the injured spinal cord. Middle lines indicate medians and markers indicate mean values. \**p* < 0.05, \*\**p* < 0.01, \*\*\**p* < 5x10^-3^, \*\*\*\**p* < 10^-3^, \*\*\*\*\**p* < 5x10^-4^, and \*\*\*\*\*\**p* < 2x10^-4^ vs. the corresponding regional values for the healthy spinal cords (Wilcoxon rank sum test). Three white matter ROIs (VP: ventral pathway, LP: lateral pathway, DP: dorsal pathway) and two gray matter ROIs (VH: ventral horn, DH: dorsal horn) are as defined in Figure 1. MD, mean diffusivity; AD, axial diffusivity; RD, radial diffusivity; FA, fractional anisotropy.

**Figure 5.**
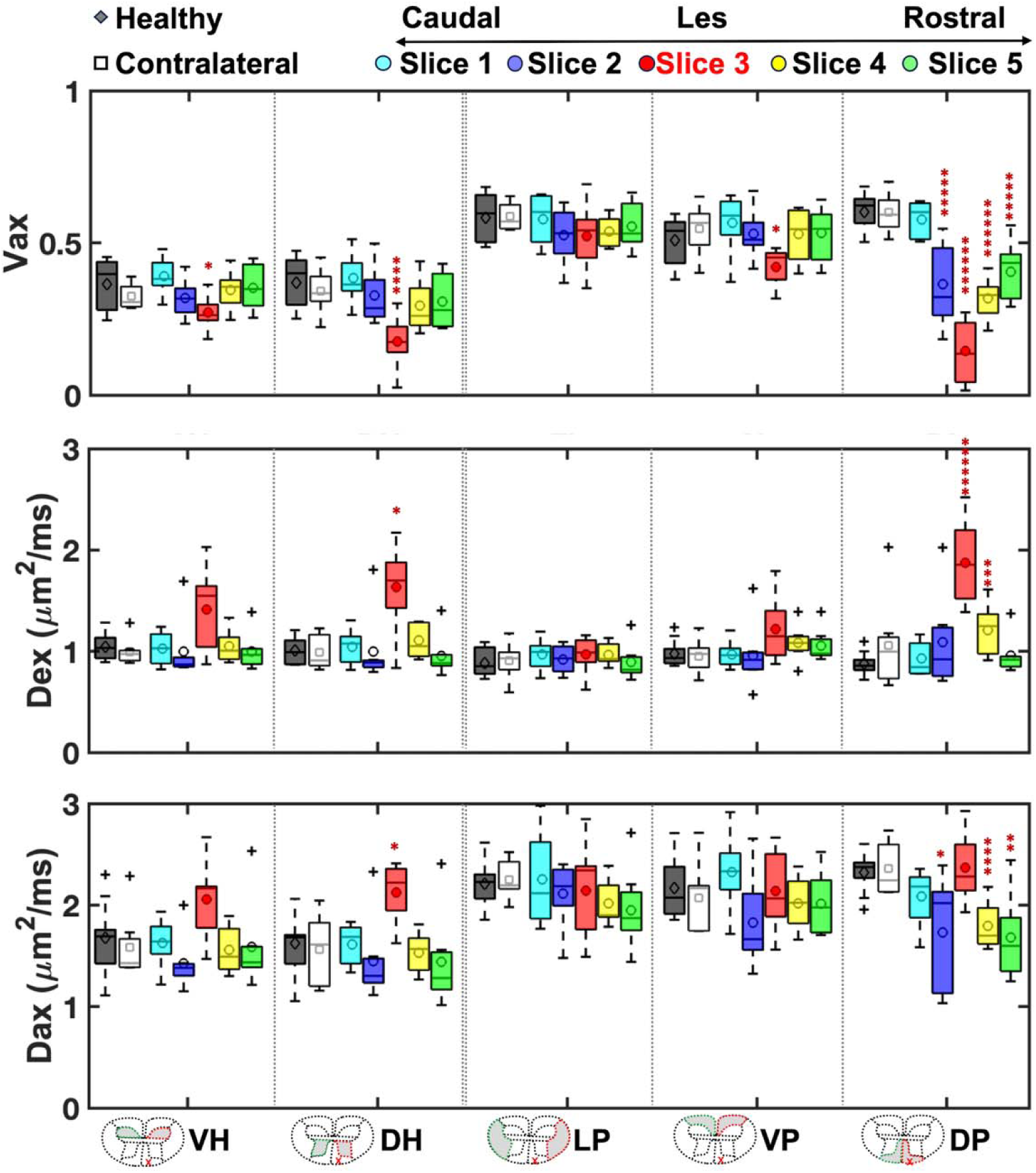
Regional comparison of SMT-derived diffusion measures in five ROIs across slices of injured spinal cords with those from healthy cords. Boxplots of SMT-derived V_ax_, D_ex_, and D_ax_ metrices show the difference for the selected ROIs from the healthy spinal cords without injury, non-lesioned contralateral side of the lesioned segment (slice 3 as indicated in Figure 3), and different segments on the lesion side from caudal to rostral (slices 1-5 as indicated in Figure 3). Middle lines indicate medians and markers indicate mean values. \**p* < 0.05, \*\**p* < 0.01, \*\*\**p* < 5x10^-3^, \*\*\*\**p* < 10^-3^, \*\*\*\*\**p* < 5x10^-4^, and \*\*\*\*\*\**p* < 2x10^-4^ vs. the corresponding regional values for the healthy spinal cords (*Wilcoxon rank sum*). Three white matter ROIs (VP: ventral pathway, LP: lateral pathway, DP: dorsal pathway) and two gray matter ROIs (VH: ventral horn, DH: dorsal horn) are as defined in Figure 1. D_ax_, intrinsic axonal diffusivity; D_ex_, extra axonal transverse diffusivity; V_ax_, apparent axonal volume fraction.

### 3.4 Tissue damage quantified by regional analysis of histological sliver staining

We also performed an ROI-based analysis of the fiber-stained tissue sections, using a histological measure HI as the ground truth to validate tissue damage detected by DTI- and SMT-derived metrics. A zoomed-in view of the ROIs in healthy spinal cord sections revealed GM and WM differences in silver staining, with WM showing higher fiber density and a more organized structure (Sup. Fig. S2). Figure 6 displays the regional quantification of HI, using the DP region as an example ROI. The selected slice showed evident axonal degeneration and demyelination in the DP region on the lesion side compared to the contralateral, non-lesioned side (Fig. 6A). From the lightness map (Fig. 6B), lightness distributions were extracted for the DP on both lesion and contralateral sides (Fig. 6C). The mean lightness across voxels in the DP on the lesion side was lower than that on the contralateral side (Fig. 6C). While HI for the DP on the lesion side showed a marked decrease compared to healthy WM, the contralateral side remained largely intact (Fig. 6C). Figure 6D compares inter- and intra-segment differences in HI across various regions and subjects. Tissue deficits were indicated by significant reductions in HI at the lesion site on the lesion side of the spinal cord, with region-specific severity classified as severe (>30% reduction), moderate (10-30%), or mild (<10%) (Fig. 6D). The extent of fiber damage showed intra-segment variation, following the pattern: DP (severe) > VP (moderate) > LP (mild) among WM regions, and DH (moderate) > VH (mild) among GM regions at the lesion level. Inter-segment differences in the DP region followed the trend: lesion center (severe) > rostral segment proximal to lesion (moderate) > caudal segment proximal to lesion (mild) > rostral segment distal to lesion (mild) > caudal segment distal to lesion (not significant) (Fig. 6D). Notably, tissue damage extended farther rostrally than caudally and showed a dorsal-to-ventral progression on the lesion side of the spinal cord (Figs. 3C and 6D). Overall, the spatial patterns of fiber density and tissue damage were consistent with regional alterations observed in diffusion MRI measures.

**Figure 6.**
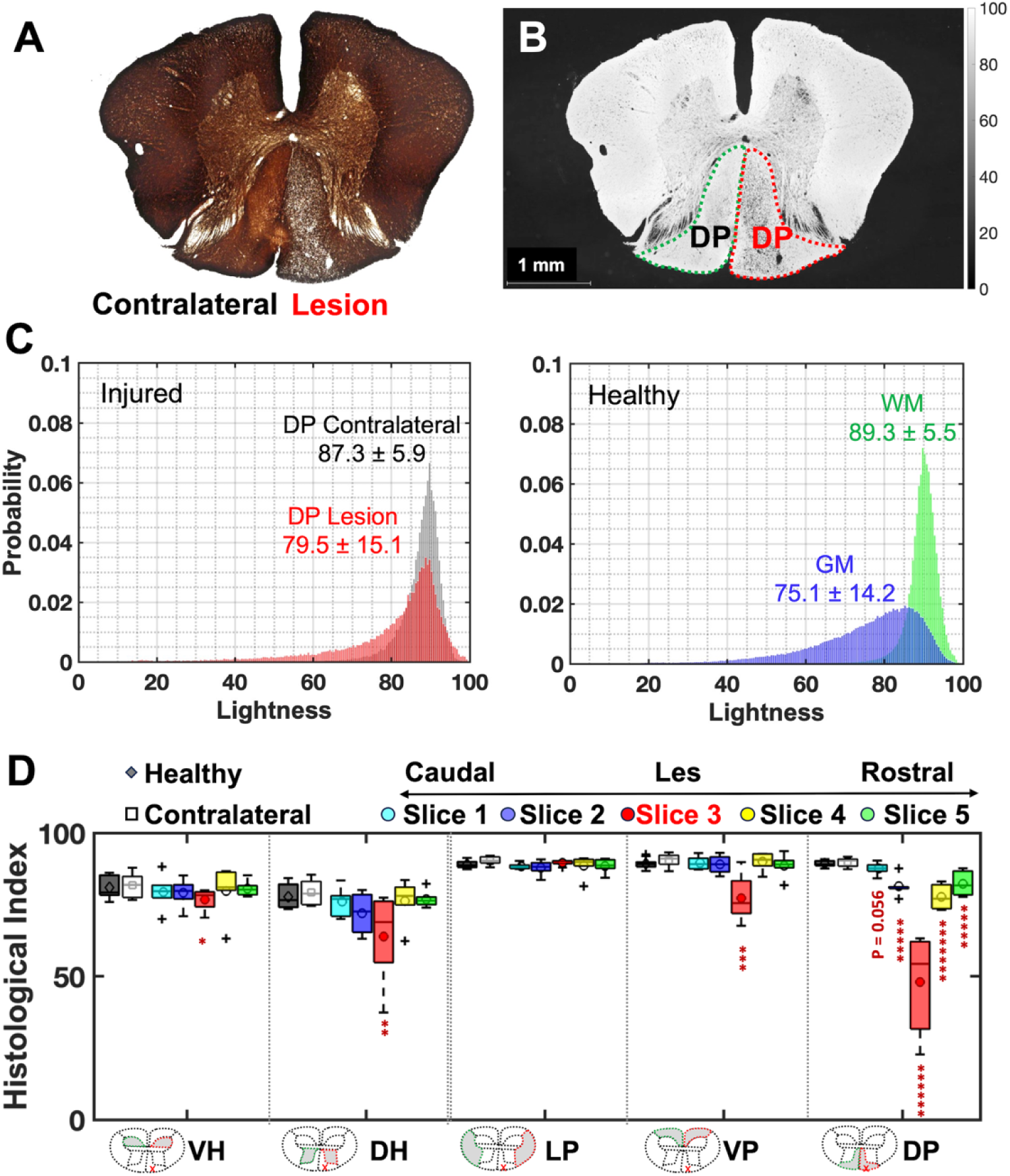
Regional histological inD_ex_ for fiber density. (A) An axial section of spinal cord tissue with silver staining. (B) Converted gray-scale map shows the lightness from 0-100. (C) Comparison of lightness distributions in dorsal pathway (DP) on non-lesioned contralateral and lesion sides in injured cord (left) and those between normal gray matter (GM) and white matter (WM) in healthy spinal cord tissue. Regional mean of lightness and respective standard deviation across voxels are provided. (D) Boxplots of regional histological inD_ex_ calculated using mean values of lightness across subjects, compare the difference for the selected ROIs from the healthy tissues without injury, non-lesioned contralateral side of the lesioned segment (slice 3 as indicated in Figure 3), and different segments on the lesion side from caudal to rostral (slices 1-5 as indicated in Figure 3). Middle lines indicate medians and markers indicate mean values. \**p* < 0.05, ** *p* < 0.01, \*\*\**p* < 5x10^-3^, \*\*\*\**p* < 10^-3^, \*\*\*\*\**p* < 5x10^-4^, and \*\*\*\*\*\**p* < 2x10^-4^ vs. the corresponding regional values for the healthy spinal cord tissues (Wilcoxon rank sum test). Three WM ROIs (VP: ventral pathway, LP: lateral pathway, DP: dorsal pathway) and two GM ROIs (VH: ventral horn, DH: dorsal horn) are as defined in Figure 1.

### 3.5 Results from ROC analysis

To determine the sensitivity and specificity of diffusion and histological metrics, we performed ROC analysis. FA demonstrated the highest TPR, TNR, and AUC for most ROIs, outperforming all other diffusion measures (Table 1 and Sup. Table S1). While the diagnostic performance of AD was low in the DP at the lesion center (AUC = 0.5), AD exhibited the large TPR, TNR, and AUC in DP for segments caudal and rostral to the lesion site, with ROC values comparable to FA (Sup. Table S1). Along the DP (Table 1), ROC values of FA and V_ax_ both followed similar spatial trend, decreasing in the following order: lesion center (slice 3) ≥ rostral segment proximal to lesion (slice 4) ≥ caudal segment proximal to lesion (slice 2) ≥ rostral segment distal to lesion (slice 5) ≥ caudal segment distal to lesion (slice 1). This was consistent with the trends observed for histological indices (Table 2).

**Table 1.**
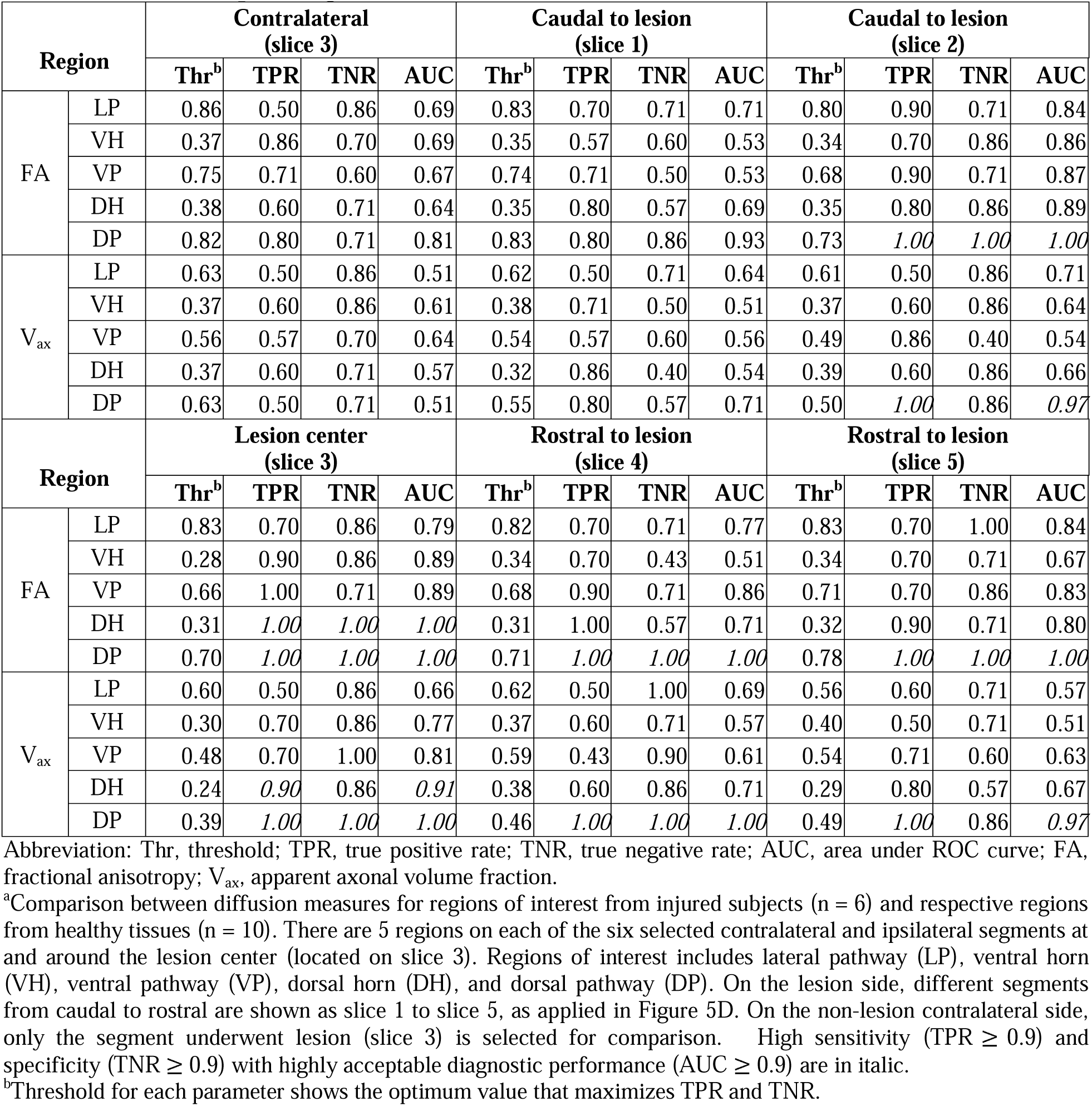
Receiver operating characteristic results of selected DTI and SMT measures^a^.

**Table 2.**
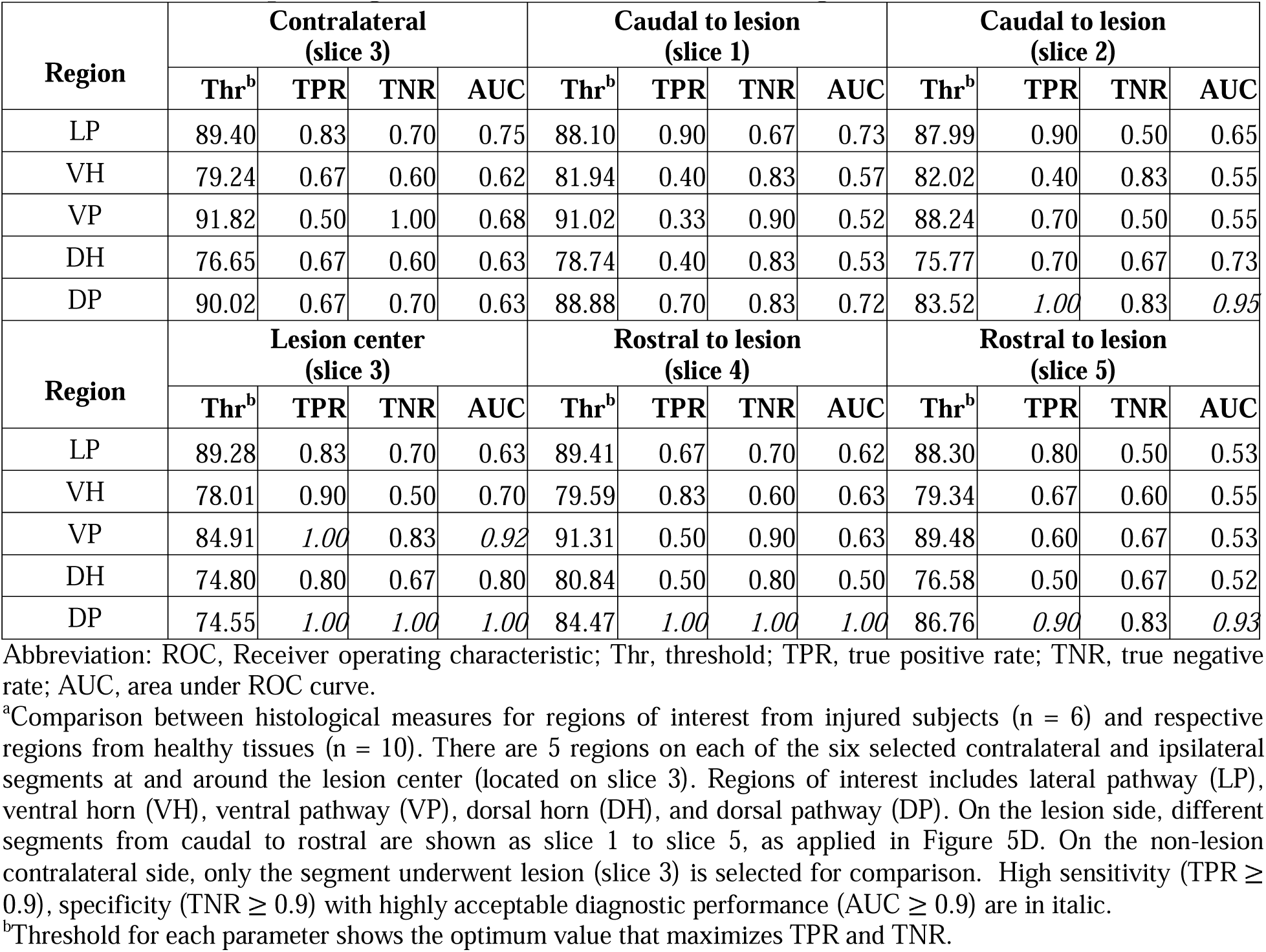
Receiver operating characteristic results of histological inD_ex_^a^.

| Region | Contralateral (slice 3) |  |  |  | Caudal to lesion (slice 1) |  |  |  | Caudal to lesion (slice 2) |  |  |  |
| --- | --- | --- | --- | --- | --- | --- | --- | --- | --- | --- | --- | --- |
|  | Thr<sup>b</sup> | TPR | TNR | AUC | Thr<sup>b</sup> | TPR | TNR | AUC | Thr<sup>b</sup> | TPR | TNR | AUC |
| LP | 89.40 | 0.83 | 0.70 | 0.75 | 88.10 | 0.90 | 0.67 | 0.73 | 87.99 | 0.90 | 0.50 | 0.65 |
| VH | 79.24 | 0.67 | 0.60 | 0.62 | 81.94 | 0.40 | 0.83 | 0.57 | 82.02 | 0.40 | 0.83 | 0.55 |
| VP | 91.82 | 0.50 | 1.00 | 0.68 | 91.02 | 0.33 | 0.90 | 0.52 | 88.24 | 0.70 | 0.50 | 0.55 |
| DH | 76.65 | 0.67 | 0.60 | 0.63 | 78.74 | 0.40 | 0.83 | 0.53 | 75.77 | 0.70 | 0.67 | 0.73 |
| DP | 90.02 | 0.67 | 0.70 | 0.63 | 88.88 | 0.70 | 0.83 | 0.72 | 83.52 | 1.00 | 0.83 | 0.95 |
| Region | Lesion center (slice 3) |  |  |  | Rostral to lesion (slice 4) |  |  |  | Rostral to lesion (slice 5) |  |  |  |
|  | Thr<sup>b</sup> | TPR | TNR | AUC | Thr<sup>b</sup> | TPR | TNR | AUC | Thr<sup>b</sup> | TPR | TNR | AUC |
| LP | 89.28 | 0.83 | 0.70 | 0.63 | 89.41 | 0.67 | 0.70 | 0.62 | 88.30 | 0.80 | 0.50 | 0.53 |
| VH | 78.01 | 0.90 | 0.50 | 0.70 | 79.59 | 0.83 | 0.60 | 0.63 | 79.34 | 0.67 | 0.60 | 0.55 |
| VP | 84.91 | 1.00 | 0.83 | 0.92 | 91.31 | 0.50 | 0.90 | 0.63 | 89.48 | 0.60 | 0.67 | 0.53 |
| DH | 74.80 | 0.80 | 0.67 | 0.80 | 80.84 | 0.50 | 0.80 | 0.50 | 76.58 | 0.50 | 0.67 | 0.52 |
| DP | 74.55 | 1.00 | 1.00 | 1.00 | 84.47 | 1.00 | 1.00 | 1.00 | 86.76 | 0.90 | 0.83 | 0.93 |

Abbreviation: ROC, Receiver operating characteristic; Thr, threshold; TPR, true positive rate; TNR, true negative rate; AUC, area under ROC curve.
<sup>a</sup>Comparison between histological measures for regions of interest from injured subjects (n = 6) and respective regions from healthy tissues (n = 10). There are 5 regions on each of the six selected contralateral and ipsilateral segments at and around the lesion center (located on slice 3). Regions of interest includes lateral pathway (LP), ventral horn (VH), ventral pathway (VP), dorsal horn (DH), and dorsal pathway (DP). On the lesion side, different segments from caudal to rostral are shown as slice 1 to slice 5, as applied in Figure 5D. On the non-lesion contralateral side, only the segment underwent lesion (slice 3) is selected for comparison. High sensitivity (TPR ≥ 0.9), specificity (TNR ≥ 0.9) with highly acceptable diagnostic performance (AUC ≥ 0.9) are in italic.
<sup>b</sup>Threshold for each parameter shows the optimum value that maximizes TPR and TNR.

### 3.6 Regional correlations between diffusion MRI and histological measures

To determine which diffusion parameter(s) best detect the spatial distribution of tissue damage, we correlated differences in diffusion metrics with corresponding HI at each ROI. The most severe tissue damage, indicated by significant reductions in histological fiber density in the DP at and around the lesion site (Fig. 6), was strongly associated with the lowest V_ax_ and FA values in the DP of the lesion center and adjacent segments (Figs. 4-5). To further assess this relationship, we examined the correlations between regional mean diffusion metrics and histological measure HI. Figure 7 shows the regional correlations across six subjects, each of whom had both diffusion and histological data for various ROIs in different segments. D_ax_ and D_ex_ demonstrated strong positive correlations with AD and RD, respectively. V_ax_ exhibited a particularly strong positive correlation with FA. Additionally, both V_ax_ and FA positively correlated with HI, whereas D_ex_, RD, and MD showed negative correlations with HI (Fig. 7). In contrast, AD and D_ax_ did not exhibit strong correlations with HI. Among all diffusion metrics, V_ax_ exhibited the strongest positive correlation with HI. Figure 7B presents the linear regression analysis between HI and V_ax_, with cyst-dominated ROIs (HI < 50) indicated by a plus symbol. When cyst-dominated ROIs were excluded, the correlation coefficient (*r* value) between V_ax_ and HI increased from 0.770 to 0.795. We also examined the correlations between regional changes in diffusion and histological measures (Fig. 8). The normalized difference in axonal volume fraction (ΔV_ax_) also showed much stronger regional correlation with the corresponding change in histological inD_ex_ (ΔHI) than did the change in fractional anisotropy (ΔFA).

**Figure 7.**
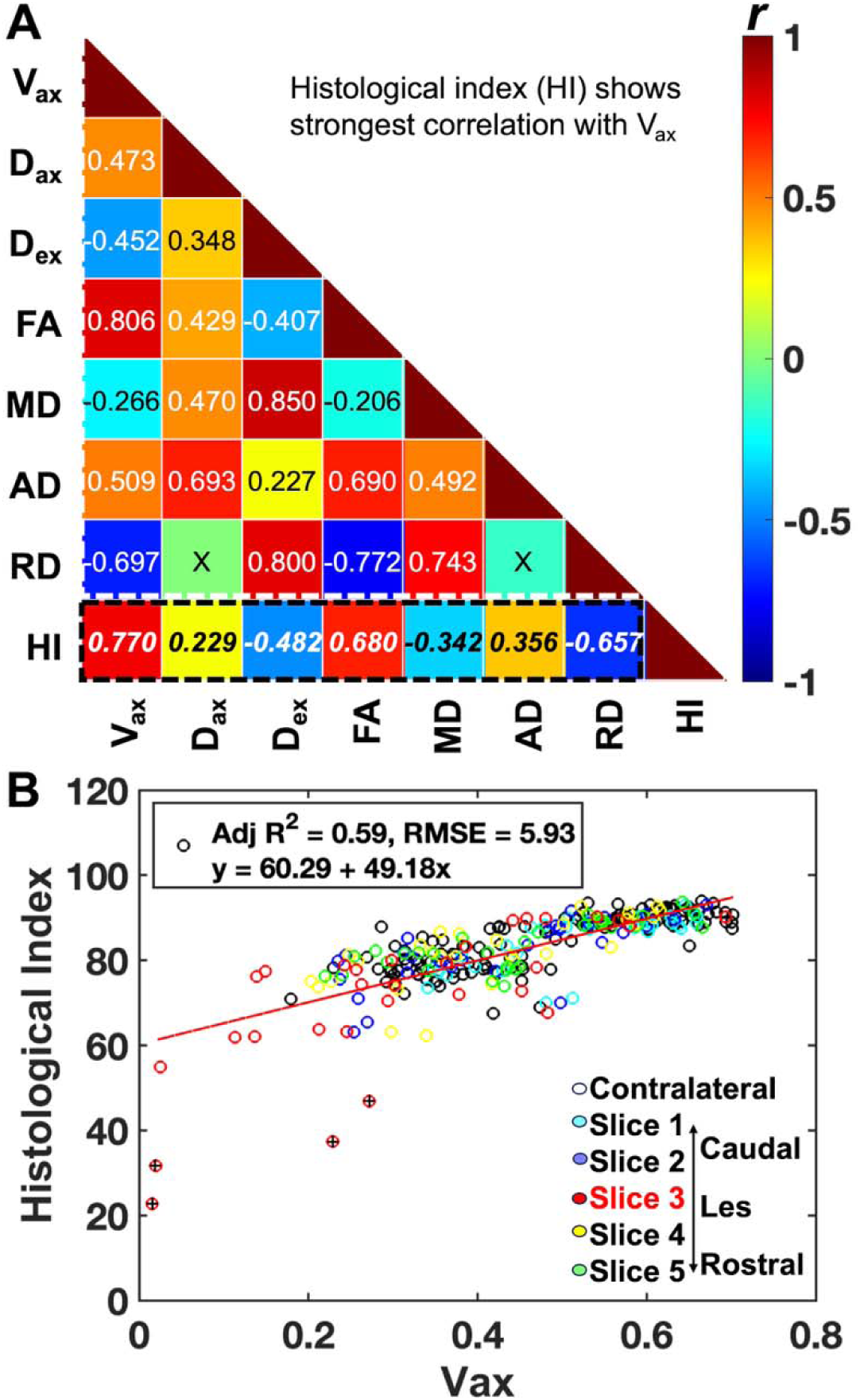
Regional correlations between diffusion and histological measures. (A) Matrix plot of correlation coefficients (*r* values) between different SMT- and DTI-derived metrics and histological inD_ex_ (HI) of fiber density. ‘x’ indicates correlation that is not significant (*p* > 0.05). Diffusion measures: V_ax_, apparent axonal volume fraction; D_ax_, intrinsic axonal diffusivity; D_ex_, extra axonal transverse diffusivity; FA, fractional anisotropy; MD, mean diffusivity; AD, axial diffusivity; RD, radial diffusivity. Number of entries per injured subject is 50 (10 ROIs on both control and lesion sides for each of the 5 slices). Total 300 entries for each metrics (number of subjects = 6) were included in correlation analysis. (B) Linear regression curve showing the relationship between histological inD_ex_ and V_ax_. The regression statistics including adjusted *R*^2^ and root mean squared error (RMSE) from modeling are included to show the goodness of predication for histological inD_ex_. The cysts were indicated by + symbols (Histological inD_ex_ < 50).

**Figure 8.**
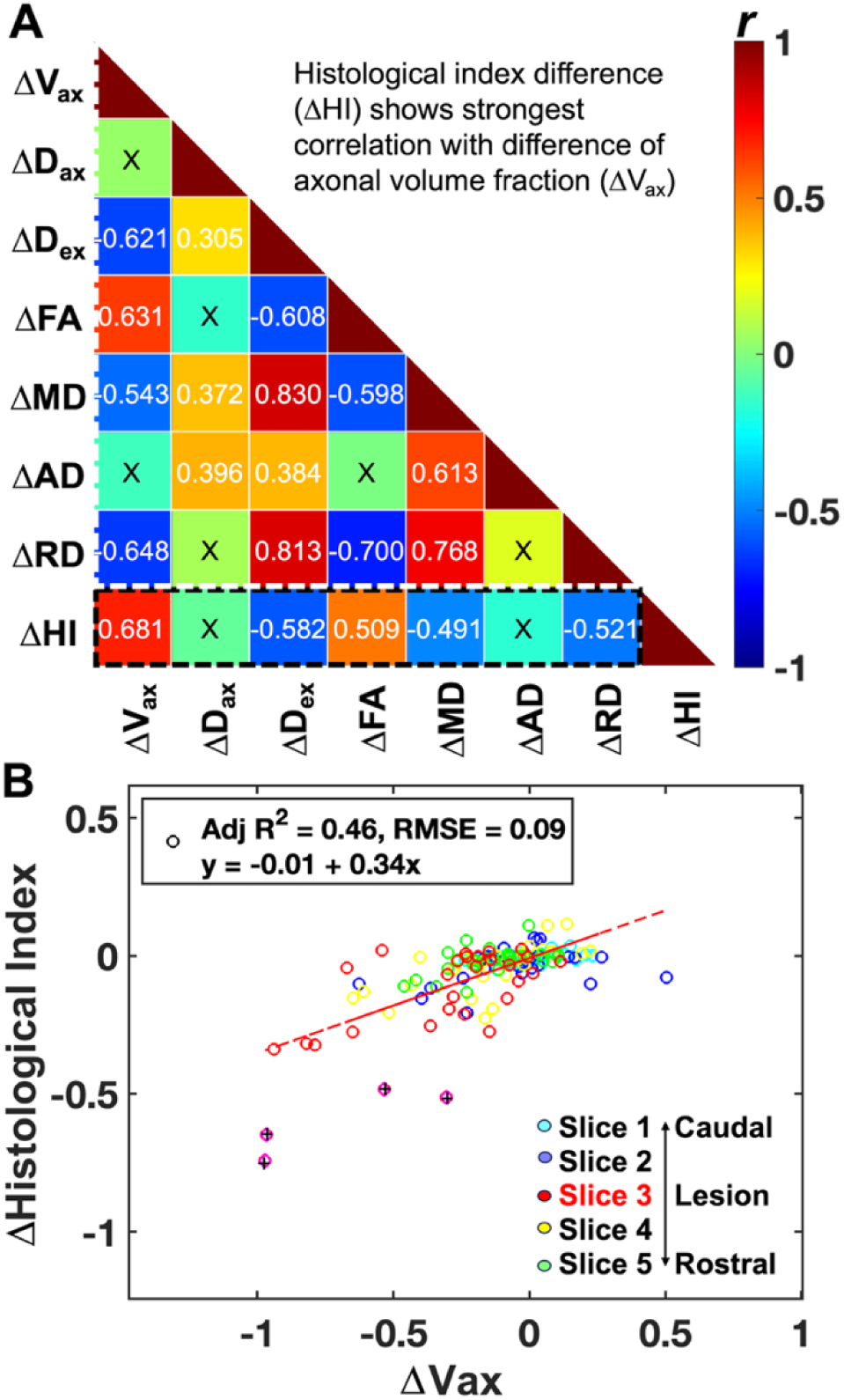
Regional correlations between normalized differences of diffusion and histological measures across all the regions within the injured spinal cord. (A) Matrix plot of correlation coefficients (*r* values) between normalized regional differences of different SMT- and DTI-derived metrics and that of histological inD_ex_ (ΔHI) of the injured spinal cords. The symbol ‘x’ indicates correlation that is not significant (*p* > 0.05). Normalized regional difference of axonal volume fraction between respective regions on the lesion and non-lesioned contralateral sides: ΔV_ax_ = (lesion-side V_ax_ – contralateral V_ax_)/contralateral V_ax_. The normalized regional differences of other diffusion metrics and histological measures were calculated similarly. ΔD_ax_, ΔD_ex_, ΔFA, ΔMD, ΔAD, and ΔRD are normalized regional differences of intrinsic axonal diffusivity, extra axonal transverse diffusivity, fractional anisotropy, mean diffusivity, axial diffusivity, and radial diffusivity, respectively. ΔHI, normalized difference of histological inD_ex_. Number of entries per injured subject is 25 (5 entries for each of the 5 slices). Total 150 entries for each metrics (number of subjects = 6) were included in correlation analysis. (B) Linear regression curve showing the relationship between ΔHistological InD_ex_ and ΔV_ax_. The regression statistics including adjusted *R*^2^ and root mean squared error (RMSE) from modeling are included to show the goodness of predication for Δhistological inD_ex_. The cysts were indicated by + symbols (Histological inD_ex_ < 50).

## 4 DISCUSSION

Our study demonstrated the technological feasibility of using the combination of DTI and SMT to detect region-specific tissue damage caused by traumatic SCI at high MRI field strength. Additionally, we have found the optimal diffusion measures FA and V_ax_ in detecting the degree and spatial spread of tissue damage across segments along the spinal cord, based on their sensitivity, specificity, and regional correlation with histological measure of fiber damage. Our results highlight the clinical translational potential of diffusion parameters FA and V_ax_ as biomarkers of fiber loss, degeneration, and demyelination caused by SCI.

### 4.1 Performance of MRI metrics from DTI and SMT for detecting region-specific fiber damage

A focal unilateral DCL initiates axonal damage that can progress within the injured segment dorsoventrally and across multiple spinal segments. Our results demonstrate that both DTI and SMT reliably detect lesion-related changes, with strong diagnostic performance (Figs. 4–5, Sup. Table S1). While most diffusion metrics are highly sensitive to severe damage at the lesion site, their ability to detect mild to moderate WM fiber degeneration, demyelination and loss of fibers in distal tissue (2 segments away from the lesion site) is particularly valuable. AD and D_ax_, though less sensitive than RD and D_ex_ at the lesion center, were more effective in detecting moderate and mild damage. Notably, AD showed the highest sensitivity, specificity, and overall performance in identifying mild to moderate axonal damage in dorsal column regions adjacent to and distal from the lesion (Sup. Table S1). FA consistently demonstrated strong diagnostic power across severity levels.

By comparing diffusion MRI metrics with histological evaluation of fiber damage, we examined which diffusion MRI metrics that spatially align closely with ground-truth histological measures. While AD and D_ax_ decreased with disrupted fibers in rostral and caudal segments, they could increase at the lesion site due to fluid accumulation and cyst formation, contributing to weaker correlations with histological measure HI (Fig. 7). In contrast, V_ax_ and FA decreased in regions of fiber damage and showed strong positive correlations with HI (Figs. 7–8). V_ax_ correlated more strongly than FA, both in absolute values (V_ax_ vs. HI) and in regional changes (ΔV_ax_ vs. ΔHI), due to its direct reflection of axonal volume. As V_ax_ represents intra-neurite or intra-axonal volume,^18^ its reduction indicates fiber damage and loss. FA, while sensitive to fiber organization, does not directly quantify fiber density and may be less reliable in complex or disorganized tissue. Overall, V_ax_ showed the strongest regional correlation with histological fiber density and best captured the spatial distribution of fiber damage (Figs. 7–8).

### 4.2 The combination of DTI and SMT detects regional characteristics following DCL

Regional histological quantification using silver staining provided a robust measure of fiber density across ROIs and confirmed the observations from diffusion MRI. HI, derived from silver staining, highlighted fiber bundles with dense staining and identified disruption, degeneration, and demyelination of axons in WM tracts and neurites in GM following SCI. Our results demonstrated strong positive correlations between diffusion metrics (particularly V_ax_ and FA) and the severity of regional damage assessed histologically (Figs. 7–8). These metrics captured clear spatial patterns of fiber damage following selective DCL in a dorsoventral order (WM: DP > VP > LP, and GM: DH > VH) and rostrocaudal trends along the dorsal column (DP: lesion center > rostral segment proximal to lesion > caudal segment proximal to lesion > rostral segment distal to lesion > caudal segment distal to lesion). The changes in segments away from the lesion are mainly WM (dorsal column) degeneration and demyelination. At and around the lesion site, tissue damage is much more complex, including GM. Different diffusion measures reflect distinct aspects of tissue microstructure, including fiber density, orientation, and composition. However, pathological factors such as demyelination, axonal loss, cysts, and gliosis can influence diffusion signals in complex ways, sometimes leading to nonlinear relationships between diffusion measures and histology. Combining multiple measures improves the accuracy of assessing SCI severity (Figs. 4–5).

Our findings underscore the value of combination of DTI and SMT as a noninvasive approach for characterizing regional damage and monitoring progression of WM fiber damage. Diffusion abnormalities were most pronounced at the lesion site and extended more prominently rostrally than caudally—consistent with known anterograde degeneration in ascending sensory pathways, as previously shown with magnetization transfer MRI.^5, 30^ While dorsal lesion damage reflects direct injury, remote tissue changes likely involve inflammation-related demyelination and gliosis.^4^ Secondary injury mechanisms such as neurotransmitter imbalance and excitotoxicity can contribute to this extended degeneration.^4, 7, 31^ The formation of cysts and regional tissue damage following SCI in this primate SCI model have been extensively discussed in previous studies.^4, 5, 30^ The ability to noninvasively detect such fiber loss in primate models has important clinical implications for early SCI diagnosis, monitoring, and evaluating therapeutic outcomes.

### 4.3 Advantages and limitations of SMT application for SCI

Applying SMT to SCI requires validating two key assumptions: that transverse diffusivity inside axons is zero, and that extra axonal transverse diffusivity can be estimated via tortuosity. Axonal loss or degeneration increase extracellular space and may reduce the degree of restriction of extracellular water. A potential issue arises because long diffusion times (typically needed for highly restricted diffusion) may confound the accuracy of the tortuosity model in cases of severe axonal loss. In cystic regions dominated by fluid, partial volume effects can bias SMT estimates of V_ax_ and D_ax_. Simulation studies have shown that SMT tends to overestimate these values when the true axonal volume fraction falls below 20%.^32^ While SMT is less accurate in cystic areas where V_ax_ < 0.3 (or HI < 0.5), it performs well across a physiological range of axonal volume fractions (V_ax_: 0.3-0.7), which encompass the physiological range of axon volume fractions (Fig. 7). Therefore, SMT remains a reliable model for assessing tissues surrounding the lesion but should be interpreted with caution in cyst-dominated regions associated with SCI.

This study supports the clinical potential of combining DTI and SMT, as both are compatible with standard MRI hardware.^18^ SMT-derived V_ax_ showed the strongest regional correlation with histological fiber damage, highlighting its value. However, SMT’s reliance on spherical mean signals across multiple *b*-values makes it more susceptible to motion, physiological noise, and B1 inhomogeneity. Consequently, although V_ax_ outperformed other metrics in correlation with fiber density, DTI-based FA still offered slightly better overall sensitivity and specificity.

## 5 CONCLUSIONS

DTI and SMT offer complementary insights into regional fiber damage following unilateral DCL, capturing both intra-segmental tissue damage in the order of DP > VP > LP and inter-segmental tissue damage in the order of lesion center > rostral > caudal. These techniques provide tissue-specific metrics related to axonal volume fraction and diffusivity. AD is more sensitive to moderate and mild fiber damage, while RD is more responsive to severe damage.

Among diffusion metrics, FA demonstrates the highest sensitivity and specificity for region-specific fiber damage across ROIs, whereas V_ax_ shows the strongest regional correlation with histologic measure of fiber density, making both reliable biomarkers of spinal cord damage. Overall, combining DTI and SMT enables a robust and comprehensive assessment of axonal pathology following SCI.

## ACKNOWLEDGEMENTS

We thank Mrs. Chaohui Tang and Mr. Fuxue Xin of the Vanderbilt University Institute of Imaging Science for their assistance in animal preparation and care during MRI data collection. We also thank Dr. Ming Lu, Mr. Ken Wilkens, and Dr. Xinqiang Yan for customizing coils for cervical spinal cord imaging. This study is supported by DOD grant W81XWH-17-1-0304, and NIH grant NS092961.

## SUPPORTING INFORMATION

Supporting Table S1. ROC results of DTI and SMT measures.

Supporting Figure S1. Representative MR images.

Supporting Figure S2. Regions of interest defined on histological section.

## Abbreviations

AD: axial diffusivity
AUC: area under the curve
D_ax_: intrinsic axonal diffusivity
D_ex_: extra axonal transverse diffusivity
DH: dorsal horn
DP: dorsal pathway
DTI: diffusion tensor imaging
FA: fractional anisotropy
GM: gray matter
HI: histological inD_ex_
LP: lateral pathway
MD: mean diffusivity
RD: radial diffusivity
ROC: receiver operating characteristic
ROI: region of interest
SCI: spinal cord injury
SMT: spherical mean technique
TNR: true negative rate
TPR: true positive rate
V_ax_: apparent axonal volume fraction
VH: ventral horn
VP: ventral pathway
WM: white matter

