## Supporting Information for "Detection of Region-specific Fiber Damage within Injured Spinal Cord Using Advanced Diffusion MRI"

**MATERIAL AND METHODS**

**SCI model: Dorsal column section**

Ten male adult squirrel monkeys (*Saimiri sciureus*) were studied, and seven of them underwent a unilateral dorsal column transection at cervical level 5 (C5). In brief, under surgical level anesthesia and aseptic conditions, the dorsal portion of the cervical spinal cord at the C4-C5 vertebrae level was exposed. The dorsal column was transected on one side using fine surgical scissors at the rostral C5 level. Each lesion was 2 mm deep, extending from the midline to the spinal nerve entering zone (approximately 2 mm in width). The dura was replaced with a small piece of gelfilm, and the wound was enclosed. Details of the surgical procedures can be found in previous publications.^1-3^

***In vivo* MRI data acquisition**

Magnetization transfer contrast (MTC) weighted structural images in different orientations were acquired with enhanced contrast for placing diffusion-weighted MRI. MTC images, with an in-plane resolution of 0.313 x 0.313 mm^2^, were acquired using a gradient echo sequence (TR/TE=220/3.24 ms, matrix size=128x128), which incorporated a Gaussian saturation pre-pulse (saturation flip angle θ_sat_ = 820°, duration = 10 ms, RF offset = 5000 Hz). Ten slices were obtained from each orthogonal imaging plane: thicknesses were 0.5 mm for coronal, 0.75 mm for sagittal, and 2 mm for axial slices.

High-resolution magnetization transfer contrast (MTC) and T_2_-weighted (T2W) anatomical images were acquired for structural comparison and ROI delineation using the same geometry as diffusion MRI. MTC images, with an in-plane resolution of 0.125 x 0.125 mm^2^, were acquired using a gradient echo sequence (TR/TE=220/4.17 ms, matrix size=256x256), which incorporated a Gaussian saturation pre-pulse (saturation flip angle θ_sat_ = 820°, duration = 10 ms, RF offset = 5000 Hz). T2W images, with an in-plane resolution of 0.250 x 0.250 mm^2^, were acquired using a fast spin echo sequence (TR/TE=3000/20 ms, matrix size=128x128).

**Histology**

One or two weeks after MRI scan, animals were euthanized with a lethal dose of sodium pentobarbital and then perfused with 4% paraformaldehyde. Post-mortem spinal cords and brainstems were removed and fixed overnight before being cut in 40-μm thick coronal or axial sections. Paraffin tissue sections were stained with several markers^4, 5^ including silver staining of myelin and fiber.^6^ The sliver staining method primarily targets specific proteins and structures within neurons, allowing for the visualization of neural components, particularly fibers including axonal structure and myelin integrity.^6^ Silver staining is sensitive in detecting and visualizing the degree and extent of fiber damage in the spinal cord, including axonal degeneration and myelin alteration.^5, 6^ In white matter (WM), silver staining highlights the myelinated axon prominently since silver has an affinity for the proteins in myelin. WM typically shows up as intensely stained areas due to the abundance of myelinated fibers. The staining in WM can help to detect disrupted and demyelinated axons within WM tracts. In gray matter (GM), silver staining highlights the neurites, including dendrites and unmyelinated axons. The staining tends to be more diffuse and less intense compared to the WM, as GM has fewer fibers.

**Data Analysis**

Due to the high prevalence rate of targeted dorsal column lesion and possible high regional sensitivity, the requested sample size for receiver operating characteristic (ROC) analysis for this injury model is small. From ROC analysis, we obtained the true positive rate (TPR), true negative rate (TNR), and area under the ROC curve (AUC).

$TPR= \frac{TP}{TP+FN}$	(3)

$TNR= \frac{TN}{TN+FP}$	(4)

Where TP represents the number of true positives, FN the number of false negatives, TN the number of true negatives, and FP the number of false positives. For each measure, the corresponding values of each region from healthy tissues were used as gold standard for comparison. High sensitivity (TPR $\geq$ 0.9) and high specificity (TNR $\geq$ 0.9) with highly acceptable diagnostic performance (AUC $\geq$ 0.9) are highlighted in Supporting Tables S1.

**REFERENCES**

1.	Qi HX, Chen LM, Kaas JH. Reorganization of somatosensory cortical areas 3b and 1 after unilateral section of dorsal columns of the spinal cord in squirrel monkeys. Research Support, N.I.H., Extramural

Research Support, Non-U.S. Gov't. *J Neurosci*. Sep 21 2011;31(38):13662-75. doi:10.1523/JNEUROSCI.2366-11.2011

2.	Chen LM, Qi HX, Kaas JH. Dynamic reorganization of digit representations in somatosensory cortex of nonhuman primates after spinal cord injury. Research Support, N.I.H., Extramural

Research Support, Non-U.S. Gov't. *J Neurosci*. Oct 17 2012;32(42):14649-63. doi:10.1523/JNEUROSCI.1841-12.2012

3.	Qi HX, Wang F, Liao CC, et al. Spatiotemporal trajectories of reactivation of somatosensory cortex by direct and secondary pathways after dorsal column lesions in squirrel monkeys. *Neuroimage*. Nov 15 2016;142:421-443. doi:10.1016/j.neuroimage.2016.08.015

4.	Wang F, Qi HX, Zu Z, et al. Multiparametric MRI reveals dynamic changes in molecular signatures of injured spinal cord in monkeys. *Magn Reson Med*. Oct 2015;74(4):1125-37. doi:10.1002/mrm.25488

5.	Wang F, Li K, Mishra A, Gochberg D, Chen LM, Gore JC. Longitudinal assessment of spinal cord injuries in nonhuman primates with quantitative magnetization transfer. *Magn Reson Med*. Apr 2016;75(4):1685-1696. doi:10.1002/mrm.25725

6.	Gallyas F. Silver staining of myelin by means of physical development. *Neurol Res*. 1979;1(2):203-9.

**Supporting Table S1. Receiver operating characteristic results of DTI and SMT measures^a^**

| ***Measure*** | | **Contralateral**  **(slice 3)** | | | | **Caudal & distal to lesion**  **(slice 1)** | | | | **Caudal &proximal to lesion**  **(slice 2)** | | | |
| --- | --- | --- | --- | --- | --- | --- | --- | --- | --- | --- | --- | --- | --- |
|  |  | ***Thr^b^*** | ***TPR*** | ***TNR*** | ***AUC*** | ***Thr^b^*** | ***TPR*** | ***TNR*** | ***AUC*** | ***Thr^b^*** | ***TPR*** | ***TNR*** | ***AUC*** |
| FA | LP | 0.86 | 0.50 | 0.86 | 0.69 | 0.83 | 0.70 | 0.71 | 0.71 | 0.80 | 0.90 | 0.71 | 0.84 |
|  | VH | 0.37 | 0.86 | 0.70 | 0.69 | 0.35 | 0.57 | 0.60 | 0.53 | 0.34 | 0.70 | 0.86 | 0.86 |
|  | VP | 0.75 | 0.71 | 0.60 | 0.67 | 0.74 | 0.71 | 0.50 | 0.53 | 0.68 | 0.90 | 0.71 | 0.87 |
|  | DH | 0.38 | 0.60 | 0.71 | 0.64 | 0.35 | 0.80 | 0.57 | 0.69 | 0.35 | 0.80 | 0.86 | 0.89 |
|  | DP | 0.82 | 0.80 | 0.71 | 0.81 | 0.83 | 0.80 | 0.86 | 0.93 | 0.73 | *1.00* | *1.00* | *1.00* |
| MD | LP | 0.89 | 0.57 | 0.80 | 0.63 | 0.80 | 0.80 | 0.71 | 0.71 | 0.85 | 0.50 | 0.71 | 0.57 |
|  | VH | 0.80 | 0.80 | 0.43 | 0.53 | 0.81 | 0.70 | 0.71 | 0.70 | 0.79 | 0.80 | 0.86 | 0.80 |
|  | VP | 0.95 | 0.50 | 0.71 | 0.53 | 0.90 | 0.71 | 0.50 | 0.57 | 0.86 | 0.70 | 0.71 | 0.74 |
|  | DH | 0.81 | 0.71 | 0.40 | 0.56 | 0.85 | 0.57 | 0.70 | 0.59 | 0.78 | 0.70 | 0.71 | 0.67 |
|  | DP | 0.94 | 0.57 | 0.70 | 0.57 | 0.91 | 0.71 | 0.60 | 0.64 | 0.87 | 0.70 | 0.71 | 0.73 |
| AD | LP | 1.97 | 0.86 | 0.70 | 0.67 | 2.07 | 0.57 | 0.80 | 0.56 | 1.95 | 0.60 | 0.86 | 0.76 |
|  | VH | 1.18 | 0.71 | 0.60 | 0.63 | 1.18 | 0.57 | 0.60 | 0.51 | 1.03 | 0.90 | 0.86 | 0.83 |
|  | VP | 1.89 | 0.86 | 0.50 | 0.63 | 1.73 | 0.90 | 0.43 | 0.56 | 1.73 | 0.90 | 0.71 | 0.81 |
|  | DH | 1.21 | 0.57 | 0.60 | 0.53 | 1.09 | 0.70 | 0.57 | 0.63 | 1.07 | 0.80 | 0.86 | 0.80 |
|  | DP | 1.96 | 0.90 | 0.57 | 0.71 | 1.97 | *0.90* | *1.00* | *0.97* | 1.80 | *1.00* | *1.00* | *1.00* |
| RD | LP | 0.28 | 0.71 | 0.60 | 0.61 | 0.34 | 0.71 | 0.90 | 0.74 | 0.28 | 0.71 | 0.60 | 0.70 |
|  | VH | 0.62 | 0.90 | 0.71 | 0.76 | 0.63 | 0.90 | 0.57 | 0.71 | 0.63 | 0.90 | 0.57 | 0.63 |
|  | VP | 0.40 | 0.80 | 0.57 | 0.67 | 0.39 | 0.80 | 0.43 | 0.50 | 0.47 | 0.43 | 0.80 | 0.53 |
|  | DH | 0.59 | 0.60 | 0.57 | 0.57 | 0.61 | 0.57 | 0.60 | 0.50 | 0.61 | 0.71 | 0.60 | 0.56 |
|  | DP | 0.34 | 0.57 | 0.90 | 0.73 | 0.34 | 0.57 | 0.90 | 0.76 | 0.38 | 0.86 | *1.00* | *0.93* |
| V_ax_ | LP | 0.63 | 0.50 | 0.86 | 0.51 | 0.62 | 0.50 | 0.71 | 0.64 | 0.61 | 0.50 | 0.86 | 0.71 |
|  | VH | 0.37 | 0.60 | 0.86 | 0.61 | 0.38 | 0.71 | 0.50 | 0.51 | 0.37 | 0.60 | 0.86 | 0.64 |
|  | VP | 0.56 | 0.57 | 0.70 | 0.64 | 0.54 | 0.57 | 0.60 | 0.56 | 0.49 | 0.86 | 0.40 | 0.54 |
|  | DH | 0.37 | 0.60 | 0.71 | 0.57 | 0.32 | 0.86 | 0.40 | 0.54 | 0.39 | 0.60 | 0.86 | 0.66 |
|  | DP | 0.63 | 0.50 | 0.71 | 0.51 | 0.55 | 0.80 | 0.57 | 0.71 | 0.50 | *1.00* | 0.86 | *0.97* |
| D_ax_ | LP | 2.11 | 0.86 | 0.30 | 0.50 | 2.14 | 0.70 | 0.57 | 0.54 | 2.20 | 0.60 | 0.57 | 0.53 |
|  | VH | 1.55 | 0.70 | 0.71 | 0.60 | 1.70 | 0.50 | 0.71 | 0.54 | 1.53 | 0.70 | 0.86 | 0.74 |
|  | VP | 1.81 | 0.80 | 0.57 | 0.67 | 2.12 | 0.86 | 0.60 | 0.66 | 1.78 | 1.00 | 0.71 | 0.79 |
|  | DH | 1.38 | 0.90 | 0.43 | 0.54 | 1.69 | 0.50 | 0.57 | 0.50 | 1.50 | 0.70 | 0.86 | 0.73 |
|  | DP | 2.52 | 0.43 | 0.90 | 0.53 | 2.25 | 0.80 | 0.71 | 0.80 | 2.20 | 0.80 | 0.86 | 0.83 |
| D_ex_ | LP | 0.90 | 0.71 | 0.70 | 0.59 | 0.92 | 0.71 | 0.70 | 0.64 | 0.91 | 0.57 | 0.70 | 0.59 |
|  | VH | 0.99 | 0.60 | 0.71 | 0.63 | 1.05 | 0.50 | 0.71 | 0.54 | 0.95 | 0.70 | 0.86 | 0.77 |
|  | VP | 0.90 | 0.70 | 0.43 | 0.53 | 0.92 | 0.71 | 0.50 | 0.50 | 0.84 | 1.00 | 0.43 | 0.61 |
|  | DH | 0.92 | 0.70 | 0.57 | 0.56 | 1.06 | 0.57 | 0.70 | 0.57 | 0.92 | 0.70 | 0.86 | 0.73 |
|  | DP | 0.96 | 0.57 | 0.80 | 0.57 | 1.02 | 0.43 | 0.90 | 0.51 | 0.92 | 0.57 | 0.80 | 0.60 |

*Continued next page*

**Supporting Table S1, Continued**

| ***Measure*** | | **Lesion Center**  **(slice 3)** | | | | **Rostral & proximal to lesion**  **(slice 4)** | | | | **Rostral & distal to lesion**  **(slice 5)** | | | |
| --- | --- | --- | --- | --- | --- | --- | --- | --- | --- | --- | --- | --- | --- |
|  |  | ***Thr^b^*** | ***TPR*** | ***TNR*** | ***AUC*** | ***Thr^b^*** | ***TPR*** | ***TNR*** | ***AUC*** | ***Thr^b^*** | ***TPR*** | ***TNR*** | ***AUC*** |
| FA | LP | 0.83 | 0.70 | 0.86 | 0.79 | 0.82 | 0.70 | 0.71 | 0.77 | 0.83 | 0.70 | 1.00 | 0.84 |
|  | VH | 0.28 | 0.90 | 0.86 | 0.89 | 0.34 | 0.70 | 0.43 | 0.51 | 0.34 | 0.70 | 0.71 | 0.67 |
|  | VP | 0.66 | 1.00 | 0.71 | 0.89 | 0.68 | 0.90 | 0.71 | 0.86 | 0.71 | 0.70 | 0.86 | 0.83 |
|  | DH | 0.31 | *1.00* | *1.00* | *1.00* | 0.31 | 1.00 | 0.57 | 0.71 | 0.32 | 0.90 | 0.71 | 0.80 |
|  | DP | 0.70 | *1.00* | *1.00* | *1.00* | 0.71 | *1.00* | *1.00* | *1.00* | 0.78 | *1.00* | *1.00* | *1.00* |
| MD | LP | 0.87 | 0.71 | 0.70 | 0.69 | 0.89 | 0.43 | 0.80 | 0.56 | 0.80 | 0.80 | 0.43 | 0.51 |
|  | VH | 0.93 | *1.00* | 0.80 | 0.94 | 0.83 | 0.71 | 0.50 | 0.56 | 0.81 | 0.70 | 0.71 | 0.73 |
|  | VP | 0.89 | 0.86 | 0.50 | 0.66 | 0.94 | 0.50 | 0.71 | 0.56 | 0.88 | 0.70 | 0.57 | 0.61 |
|  | DH | 0.96 | *1.00* | *0.90* | *0.99* | 0.85 | 0.86 | 0.70 | 0.79 | 0.80 | 0.70 | 0.71 | 0.69 |
|  | DP | 1.15 | 0.86 | *1.00* | *0.96* | 1.01 | 0.43 | 0.90 | 0.56 | 0.85 | 0.80 | 0.86 | 0.79 |
| AD | LP | 1.97 | 0.50 | 0.57 | 0.50 | 1.84 | 1.00 | 0.57 | 0.66 | 1.97 | 0.57 | 0.70 | 0.51 |
|  | VH | 1.27 | 0.71 | 0.70 | 0.64 | 1.11 | 0.80 | 0.43 | 0.53 | 0.97 | 1.00 | 0.43 | 0.60 |
|  | VP | 1.87 | 0.60 | 0.86 | 0.76 | 1.67 | 1.00 | 0.71 | 0.74 | 1.81 | 0.80 | 0.71 | 0.76 |
|  | DH | 1.57 | 0.71 | 0.90 | 0.77 | 1.20 | 0.71 | 0.60 | 0.57 | 1.05 | 0.90 | 0.43 | 0.60 |
|  | DP | 2.01 | 0.70 | 0.43 | 0.50 | 1.81 | *1.00* | *1.00* | *1.00* | 1.77 | *1.00* | *1.00* | *1.00* |
| RD | LP | 0.29 | 0.71 | 0.70 | 0.73 | 0.30 | 0.71 | 0.70 | 0.73 | 0.30 | 0.86 | 0.70 | 0.80 |
|  | VH | 0.74 | 0.86 | 0.90 | 0.91 | 0.71 | 0.57 | 0.80 | 0.56 | 0.64 | 0.80 | 0.57 | 0.66 |
|  | VP | 0.55 | 0.57 | 0.90 | 0.70 | 0.45 | 0.86 | 0.70 | 0.74 | 0.48 | 0.57 | 0.80 | 0.64 |
|  | DH | 0.89 | 0.86 | 1.00 | 0.94 | 0.71 | 0.71 | 0.90 | 0.87 | 0.62 | 0.71 | 0.60 | 0.63 |
|  | DP | 0.46 | *1.00* | *1.00* | *1.00* | 0.42 | *1.00* | *1.00* | *1.00* | 0.39 | 0.86 | *1.00* | *0.94* |
| V_ax_ | LP | 0.60 | 0.50 | 0.86 | 0.66 | 0.62 | 0.50 | 1.00 | 0.69 | 0.56 | 0.60 | 0.71 | 0.57 |
|  | VH | 0.30 | 0.70 | 0.86 | 0.77 | 0.37 | 0.60 | 0.71 | 0.57 | 0.40 | 0.50 | 0.71 | 0.51 |
|  | VP | 0.48 | 0.70 | 1.00 | 0.81 | 0.59 | 0.43 | 0.90 | 0.61 | 0.54 | 0.71 | 0.60 | 0.63 |
|  | DH | 0.24 | *0.90* | 0.86 | *0.91* | 0.38 | 0.60 | 0.86 | 0.71 | 0.29 | 0.80 | 0.57 | 0.67 |
|  | DP | 0.39 | *1.00* | *1.00* | *1.00* | 0.46 | *1.00* | *1.00* | *1.00* | 0.49 | *1.00* | 0.86 | *0.97* |
| D_ax_ | LP | 1.84 | 1.00 | 0.43 | 0.51 | 2.05 | 0.80 | 0.71 | 0.73 | 1.92 | 0.90 | 0.71 | 0.77 |
|  | VH | 1.90 | 0.71 | 0.80 | 0.76 | 1.65 | 0.60 | 0.71 | 0.59 | 1.61 | 0.70 | 0.86 | 0.66 |
|  | VP | 2.08 | 0.50 | 0.57 | 0.51 | 1.85 | 1.00 | 0.43 | 0.66 | 1.98 | 0.70 | 0.57 | 0.66 |
|  | DH | 1.83 | 0.86 | 0.80 | 0.86 | 1.64 | 0.60 | 0.71 | 0.60 | 1.48 | 0.70 | 0.71 | 0.71 |
|  | DP | 2.52 | 0.43 | 0.90 | 0.53 | 2.04 | *0.90* | 0.86 | *0.96* | 1.92 | 1.00 | 0.86 | 0.89 |
| D_ex_ | LP | 0.92 | 0.71 | 0.70 | 0.70 | 0.90 | 0.86 | 0.70 | 0.74 | 0.89 | 0.43 | 0.70 | 0.50 |
|  | VH | 1.16 | 0.71 | 0.80 | 0.74 | 1.00 | 0.71 | 0.50 | 0.50 | 1.04 | 0.50 | 0.86 | 0.66 |
|  | VP | 1.04 | 0.71 | 0.80 | 0.76 | 0.99 | 0.86 | 0.70 | 0.71 | 0.96 | 0.86 | 0.60 | 0.69 |
|  | DH | 1.27 | 0.86 | 1.00 | 0.86 | 1.01 | 0.71 | 0.60 | 0.70 | 0.97 | 0.60 | 0.86 | 0.70 |
|  | DP | 1.24 | *1.00* | *1.00* | *1.00* | 0.94 | 0.86 | 0.80 | 0.91 | 0.92 | 0.57 | 0.80 | 0.66 |

Abbreviation: Thr, threshold; TPR, true positive rate; TNR, true negative rate; AUC, area under ROC curve; FA, fractional anisotropy; AD, axial diffusivity; RD, radial diffusivity; MD, mean diffusivity. V_ax_, apparent axonal volume fraction; D_ax_, intrinsic axonal diffusivity; D_ex_, extra‐axonal transverse diffusivity.

^a^Comparison between diffusion measure for regions of interest from injured subjects (n = 6) and respective regions from healthy tissues (n = 10). There are 5 regions on each of the six selected contralateral and ipsilateral segments at and around the lesion center (located on slice 3). Regions of interest includes lateral pathway (LP), ventral horn (VH), ventral pathway (VP), dorsal horn (DH), and dorsal pathway (DP). Different segments from caudal to rostral are shown as slice 1 to slice 5. Location and side of lesion are highlighted in color as applied in Figures 5-6 in the main manuscript. High sensitivity (TPR $\geq$ 0.9) and specificity (TNR $\geq$ 0.9) with highly acceptable diagnostic performance (AUC $\geq$ 0.9) are in italic.

^b^Threshold for each parameter shows the optimum value that maximizes TPR and TNR.

**Supporting Figure S1**. Comparison of representative MR images with magnetization transfer contrast (MTC), T_2_-weighted (T2W) image, non-diffusion-weighted SE-EPI (A_0_) image, diffusion-weighted SE-EPI images (A_1_-A_30_), and fractional anisotropy (FA) map.

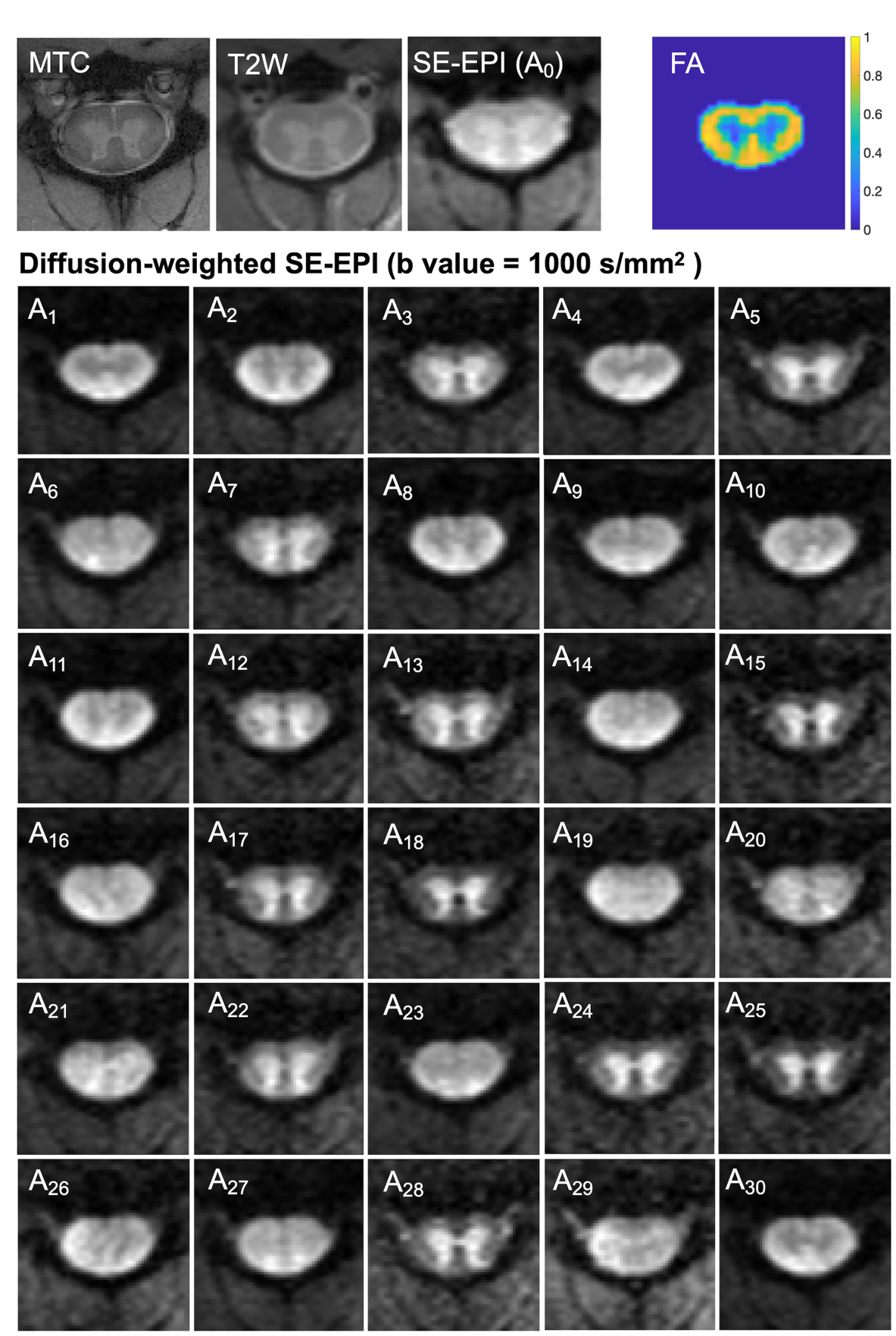

**Supporting Figure S2**. Regions of interest (ROIs) defined on characteristic histological silver staining section. (A) ROI selection for white matter regions such as ventral pathway (VP), lateral pathway (LP), and dorsal pathway (DP), and gray matter regions such as dorsal horn (DH) and ventral horn (VH). The boundaries are indicated by the dashed lines. (B-E) Enlarged ROIs showing the regional difference of fiber density. One normal cervical spinal cord section (in 40 mm thickness) was shown.

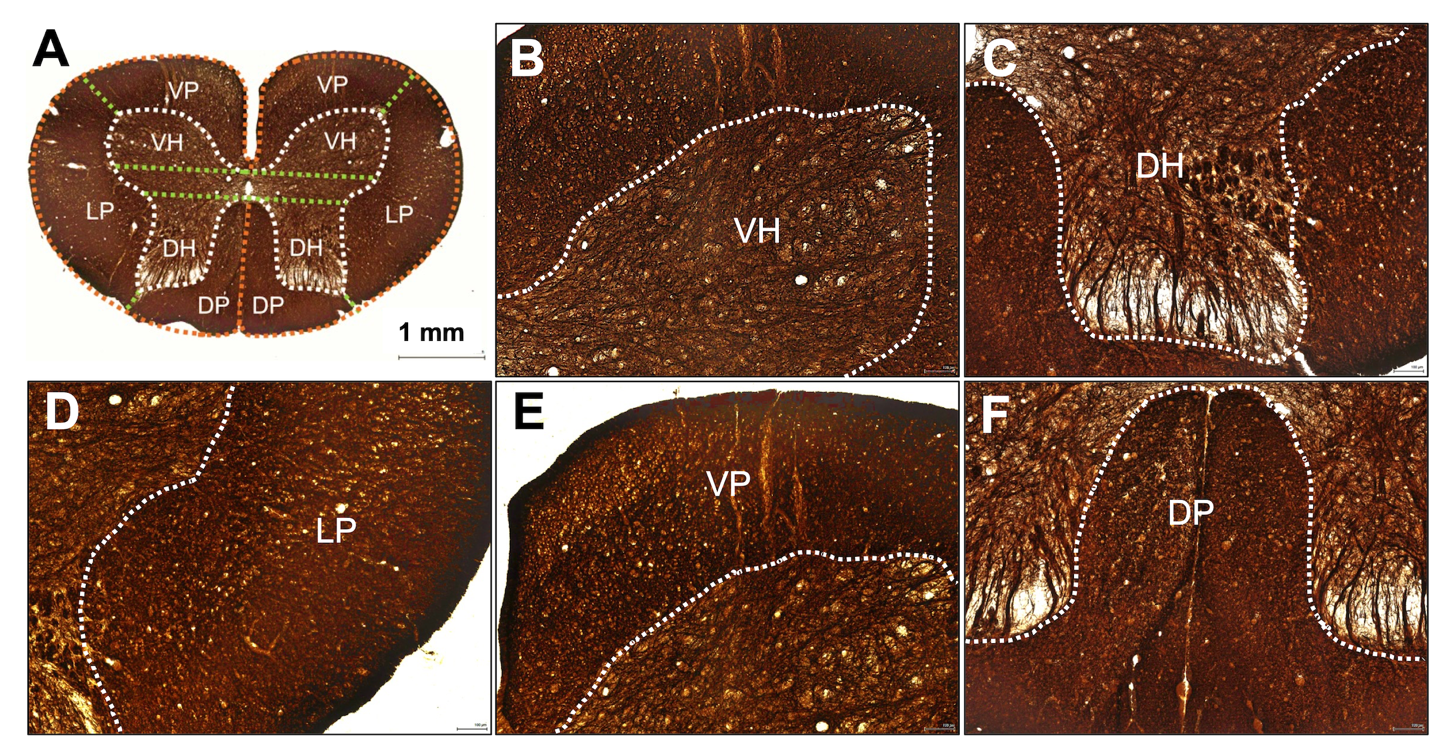
